# Multiomics plasticity in seed traits of pan-genome wheat cultivars

**DOI:** 10.1101/2024.12.20.629680

**Authors:** Utpal Bose, Jana Barbro Winkler, Elisa Sorg, Shahida A. Mitu, Gregor Huber, Robert Koller, David J. Beale, Amanda L. Dawson, Sophia Escobar-Correas, Bhabananda Biswas, Mohammad M. Rahman, Sally Stockwell, Keren Byrne, James Broadbent, Manjusha Neerukonda, Franz Buegger, Alexandros Sigalas, Klaus F. X. Mayer, Detlef Schuppan, Curtis Pozniak, Michelle L. Colgrave, Manuel Spannagl, Angéla Juhász, Jörg-Peter Schnitzler

## Abstract

The molecular basis of cultivar-level variations in polyploid wheat that enables environmental adaptation while maintaining yield and quality in polyploid wheat remains poorly understood. We conducted a detailed phenotypic assessment and multiomics analysis of nine pan-genome polyploid wheat cultivars grown under control and drought conditions. We aimed to investigate the subgenome-level variations, cultivar differences and biochemical mechanisms affecting plant fitness under moderate drought stress. Intrinsic water use efficiency, grain yield, and grain protein content and quality differed among cultivars, supporting the plasticity of drought stress responses. Biased proteome and metabolome abundance changes in response to moderate drought stress during the vegetative stage indicate different strategies for the utilization of homeologous protein isoforms assigned to the A, B, and D subgenomes. Drought effects were detected at the protein level, but significant changes were observed in central carbon pathway metabolites and micronutrient profiles. The subgenomic localization of seed storage proteins highlight differences in nutrient reservoir accumulation and emphasizes the enhanced role of S-rich prolamins in the stress response. Subgenomic variations define cultivar phenotypes by producing molecules that accumulate and enable the underlying trade-offs between environmental adaptation and yield- or quality-related traits. These variations can be used to select crops with increased stress resistance without compromising yield.

## Introduction

Wheat is one of the most commonly cultivated crops worldwide, with a production area of over 200 million hectares and an annual production of 792.9 million metric tons (*1*). Hexaploid wheat (AABBDD) provides nutritional and health benefits to approximately one-third of the global population (*2*). To feed the growing world population, wheat yields need to increase by 50% in the Year 2050 (*3*), despite the often-adverse effects of climate change on yield and yield stability (*4*). Thus, developing climate-resilient and high-yielding wheat varieties by utilizing cultivar-level variations is important for increasing productivity and stress resilience (*5*). The aim of our study was to combine comprehensive phenotyping assessments and multiomics layers to better understand the plasticity among wheat cultivars. We used wheat pan-genome cultivars from the 10+ Wheat Genome Project (*6*) to conduct a controlled drought stress experiment. We analyzed plant and seed phenotypes, grain yield, intrinsic water use efficiency, and molecular changes in the grain proteome, metabolome, and micronutrient compositions to elucidate the underlying mechanisms involved in species-level variations in response to moderate drought stress.

The bread wheat genome is approximately five times larger than the human and maize genomes, with a size of 16 Gb (*6*). The high repeat content and presence of conserved triads (homelogous genes present in all three bread wheat subgenomes) introduce complexities in the genomic analysis of wheat. Nevertheless, owing to significant technological advances, both high-quality reference genomes (*7*) and pan-genomes (*6*, *8*) have been successfully generated for bread wheat. The pan-genome resources established by the 10+ Wheat Genome Project have proven invaluable in identifying and investigating candidate genes, structural variations and allelic diversity. A comprehensive wheat pan-transcriptome atlas and analysis approach have been recently developed. These data reveal widespread cultivar-specific variation in expression patterns and gene content (*9*). Our study extends the wheat pan-genome framework to encompass the pan-proteome and pan-metabolome of seeds, with a focus on genetic variability and the dynamic changes introduced by cultivar-level variation under moderate drought stress and control conditions.

One of the most significant impacts of regional climate change is drought stress (*10*), which negatively impacts plant development and perturbs cellular processes, ultimately reducing crop productivity. Wheat grain quality and yield are closely linked to physiological processes during the vegetative and grain-filling stages. However, many current efforts address only crop genetics and focus on extreme drought, which can lead to crop failure (*11*), reduction in grain quality and changes in the content and composition of antinutritional proteins, thereby threatening food security (*12–14*). Several traits influence the ability to adapt to water scarcity, with varying degrees of impact (*15*). Extreme drought during germination or anthesis has a significant effect on yield, whereas water shortage during grain filling mainly affects grain protein quality (*14, 16*). Moderate stress, on the other hand, triggers adaptive responses that can repair stress-related damage, restore cellular homeostasis, and adapt growth to appropriate stress conditions (*17*).

In the present study, a comprehensive set of phenotypic traits, proteomes, central carbon pathway metabolites and micronutrient profiles in the grains of nine pan-genome wheat lines grown under normal and drought conditions were established and analyzed. The analysis yielded valuable insights into the wheat grain ‘omics’ and phenotypic plasticity and dynamics, including impacts on protein accumulation and distribution at the chromosome level of the three subgenomes. Additionally, cultivar-specific differences and the effects of moderate drought on grain metabolite and micronutrient composition were revealed. Furthermore, the results provide comprehensive resources that will facilitate the improvement of wheat varieties. The insights into the metabolic adaptability and proteome plasticity gained from this study will prove invaluable in future wheat breeding and grain quality research.

## Results

### Multiomics analysis of the reference pan-wheat grains establishes a novel resource for wheat research

#### Grain core/shell/cloud peptidome

We conducted a multiomics analysis of the grains from nine bread wheat cultivars (Fig. 1A, 1B) selected from the 10+ Wheat Genome Project (*6*). To investigate the global variation in the subgenome-specific proteoforms, both among the wheat subgenomes and between cultivars, we analyzed discovery proteomics data at peptide levels collected from the control samples. We detected a total of 27,386 peptides across the nine cultivars. The number of peptides identified in individual cultivars ranged between 18,006 (Weebill) and 19,587 (Chinese Spring), indicating a low level of variance in the number of detected peptides. Approximately 55.6% of the peptides were shared between cultivars representing the core peptidome; 24% were detected in at least two cultivars (shell peptidome); and 20.3% were cultivar-specific peptides (cloud peptidome) (Fig. 1C). Differences in the core and variable (shell and cloud) peptidomes revealed highly conserved grain proteome compositions. The variable peptidome was associated with defense response, oxidative stress and allocation of N-containing nutrient reservoirs. Moreover, major biological processes and enzymes involved in energy metabolism, carbohydrate metabolism, translation, protein folding and intracellular protein transport are driven by more conserved proteins (Fig. 1C, fig. S1A, fig. S1B).

**Fig. 1.**
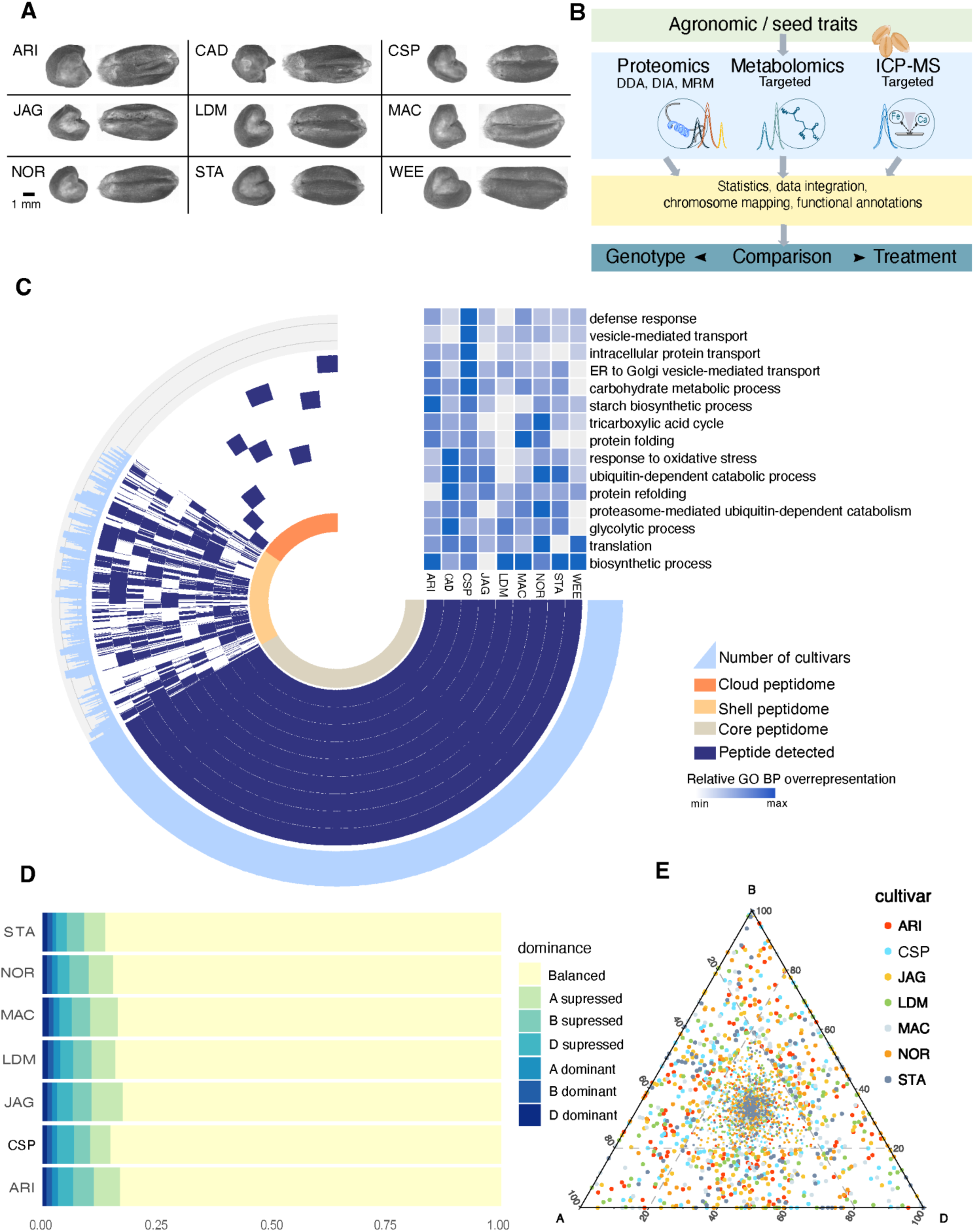
Pan-genome wheat grains reveal differences in grain proteomes. (**A**) 3D images of representative mature grains from the nine wheat cultivars. (**B**) Schematic overview of the multiomics analysis workflow. (**C**) LC-MS-based pan-peptidome level analysis showing differences in core, shell and cloud grain proteomes identified in the cultivars. (**D**) The subgenome dominance analysis shows over 80% of the peptides have balanced abundance levels between the cultivars. (**E**) Triad analysis of peptides detected from all three subgenomes. Ternary plot showing the relative abundance of triad-derived proteoforms representing the grains from nine wheat cultivars. Each dot represents a single proteoform. Triad proteoform analysis reveals balanced abundance patterns across the three subgenomes. Abbreviation of the cultivars: ARI - Arina*LrFor*, CAD - Cadenza, CSP - Chinese Spring, JAG - Jagger, LDM - CDC Landmark, MAC – Mace, NOR – Norin 61, STA – CDC Stanley, WEE – Weebill. (**D**- **E**) Only cultivars with chromosome-scale assembly are visualized.

We subsequently used quantitative targeted proteomics assays to monitor gluten proteins with the importance in end-use quality and known antinutritive characteristics (*14*). Additionally, amylase trypsin inhibitors (ATIs) that activate toll-like receptor 4 (TLR4) triggering inflammatory non-coeliac wheat sensitivity (NCWS), celiac disease (CD) (*18*) and are known wheat allergens (WA) in baker’s asthma (*19*) were measured.

A total of 136 tryptic and chymotryptic peptides specific to the major gluten protein subtypes (α-, γ-, ο-gliadins, and high-molecular-weight [HMW] and low-molecular weight [LMW] glutenins) were selected for statistical analysis (Fig. 2A). We observed significant abundance differences of gluten proteins in the cultivars Arina*LrFor* and Cadenza compared with the other cultivars at the level of each glutenin and gliadin locus encoded on chromosomes 1 and 6 groups (Fig. 2A, table S1). The abundance level differences in S-rich (α- and γ-gliadins, and LMW glutenins) and S-poor (ο-gliadins and HMW glutenins) gluten proteins indicate that storage protein accumulation is highly variable at the subgenomic and chromosome levels (Fig. 2B, fig. S3, table S1), suggesting possible differences in storage protein accumulation pathways with a direct impact on end-use quality and immunoreactive protein abundance.

**Fig. 2.**
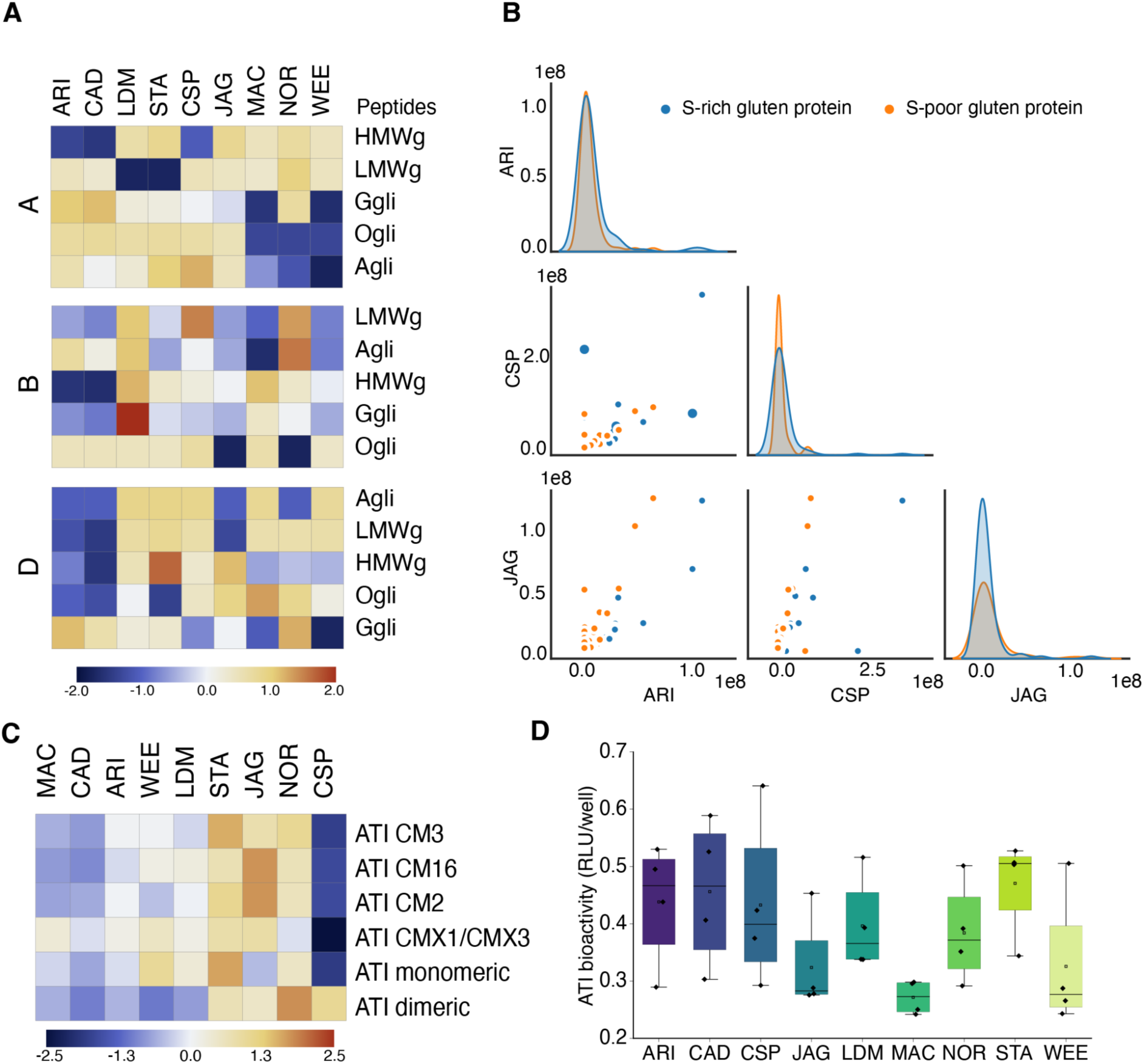
Targeted protein measurements reveal cultivar-specific differences in gluten protein and ATI profiles. (**A**) Patterns of subgenome specific gluten protein subtype abundance profiles including main gluten protein types (Agli: α-gliadin, Ggli: γ-gliadin, Ogli: -gliadin, HMWg: HMW glutenin, LMWg: LMW glutenin). (**B**) Representative abundance profiles of cultivars with diverging S-rich and S-poor gluten proteins. (**C**) Significant differences in ATI (α-amylase/trypsin inhibitor) subtype abundances across nine cultivars. ATI subtype profiles indicate differences in monomeric, dimeric and CM-type ATI levels. (**D**) Cell-based dual reporter bioassay measuring cultivar-specific TLR4 activating cellular responses to ATIs that were quantitatively extracted from 1g of respective flours. RLU, relative luminescence units.

We noted cultivar-specific differences in the abundance of ATI subtypes (20, *21*), with CM subtypes being low in the Chinese Spring, Mace, Cadenza and Arina*LrFor* cultivars and high in Jagger and CDC Stanley (Fig. 2C, table S2). The higher abundance of ATI CM3 in CDC Stanley, Norin 61 and Jagger may lead to an increased innate immune response in patients with NCWS. In addition to the ATI subtypes CM2, CM3 and CM16 subclasses, the monomeric and dimeric ATI subclasses are associated with respiratory allergies (*22*). In our results, the dimeric ATI subtype was significantly more abundant in Norin 61, whereas the monomeric subclass was significantly higher in Stanley (Fig. 2C, table S2). Finally, we measured ATI bioactivities using a dual reporter TLR4 cell line isolated from NCWS patients and detected no significant differences between cultivars (p<0.72; one-way analysis of variance [ANOVA] followed by Tukey’s multiple comparisons), corroborating our findings of unchanged ATI content in the wheat population over 100 years (*23*) (Fig. 2D).

The use of translated pan genomic gene models resulted in the capture of quantitative differences at the peptide and proteoform levels (Fig. 1C, 1D, and 1E) with increased abundances of proteins involved in DNA replication and protein synthesis in CDC Stanley, CDC Landmark, Chinese Spring, Jagger, and Weebill (Fig. 3A). The opposite relationship was observed for the remaining cultivars, namely, Arina*LrFor,* Cadenza, Norin 61, and Mace, which showed differences in proteins involved in energy metabolism and carbon partitioning.

**Fig. 3.**
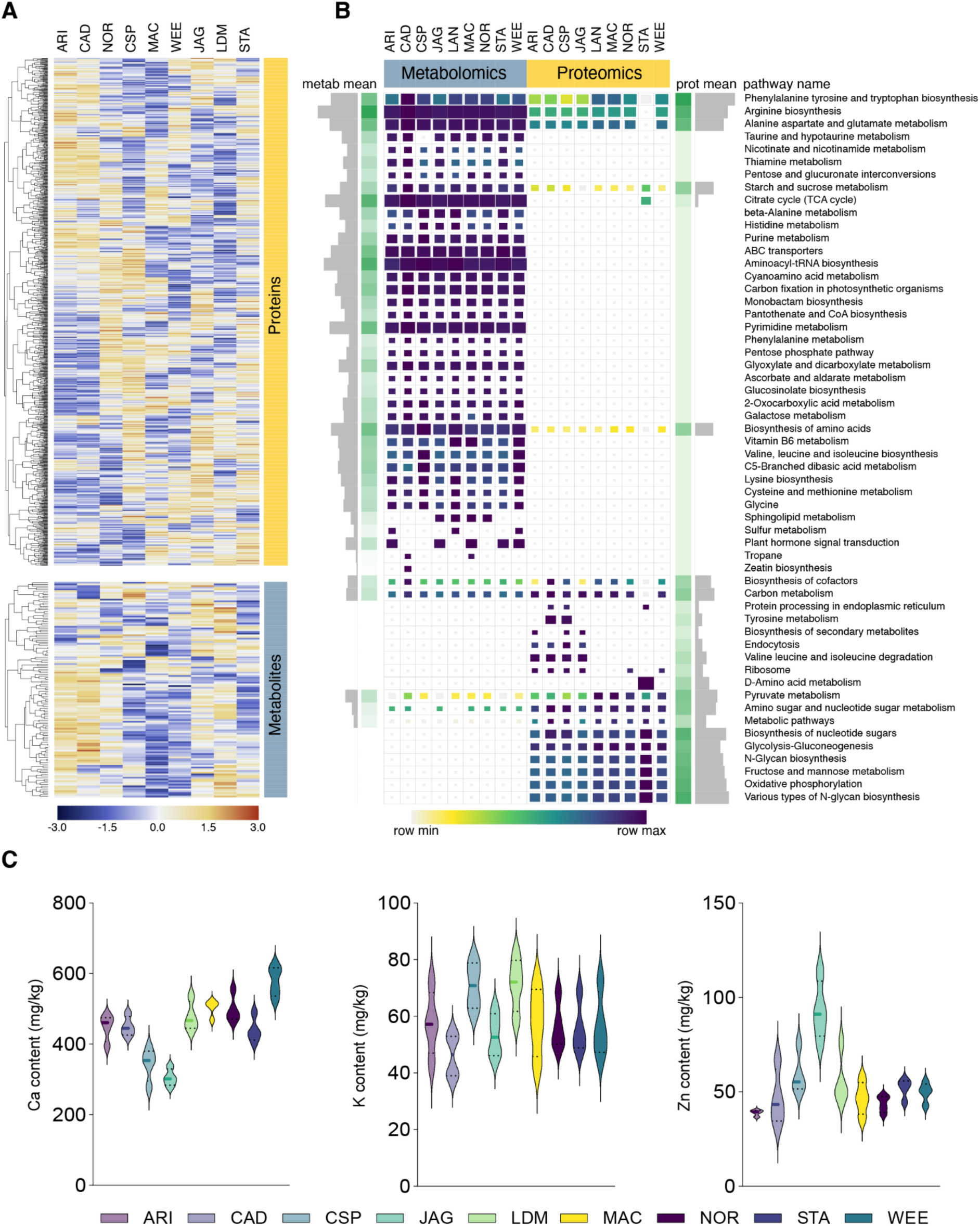
Cultivar comparisons show significant variation in protein, metabolite and micronutrient profiles. (**A**) Protein and metabolite profiles reveal genetic variability between cultivars. (**B**) Comparison of enriched KEGG pathways in the proteomics and metabolomics datasets. Proteins and metabolites representing significant differences between any pair of cultivars were used for the KEGG pathway overrepresentation analysis. Enrichment scores of significantly enriched (adj. p-value <0.05) KEGG pathways were visualized. Proteomics and metabolomics results were scaled separately. Mean enrichment scores of metabolomics and proteomics-related pathways are shown in the bar charts. (**C**) Micronutrient (Ca, K and Zn) concentrations of nine cultivars in control condition.

#### Wheat cultivars show high variability in their grain metabolic and micronutrient profiles

Central carbon metabolism metabolite and micronutrient accumulation in grains further indicated phenotypic plasticity at the cultivar level (Fig. 3A). A total of 127 metabolites were significantly different (adjusted p<0.05) between the cultivars (Fig. 3A, fig. S4), mostly metabolites involved in carbohydrate, amino acid and energy metabolism (Fig. 3B); the differentially abundant primary metabolites were also overrepresented in these six pathways (Fig. 3B, table S3).

Finally, the comparison of grain micronutrients revealed that nine micronutrients differed in abundance between cultivars (adj p<0.05) (table S4). We compared the differences in the concentrations of three main micronutrients: calcium (Ca), potassium (K) and zinc (Zn) among the cultivars (Fig. 3C). These micronutrients were selected because they play a key role in micronutrient deficiency in the developing world, which is dependent mainly on cereal-based diets (*24*). The highest Ca concentration was detected in Weebill, whereas the highest Zn concentration was measured in Jagger, which was >2-fold higher than that detected in Arina*LrFor*. The highest K concentration was detected in Arina*LrFor*, while an approximately 49% lower concentration was detected in Jagger. These results provide clear evidence of variation in micronutrient accumulation in wheat grains at the cultivar level, which is consistent with recent findings on quantitative trait locus (QTL)-level variations in six minerals across eleven wheat cultivars (*25*).

### Phenotypic variability of pan-genome wheat cultivars under moderate drought stress

#### Water restriction affects plant morphological traits

We conducted a controlled drought tolerance experiment on the nine wheat cultivars (*6*). Moderate drought stress was applied 62 days after sowing (DAS), when the plants were in the stem elongation stage (BBCH 32/33) (*26*) (fig. S5, S6, S7). Under moderate drought conditions, cumulative irrigation over the entire growing season averaged 10 L for all cultivars, which was 59% of that under the control condition (17 L) (fig. S7B).

The dynamics of plant growth were assessed with digital biomass (m³) as the key performance indicator (Fig. 4A, fig. S8A). After the onset of drought treatment at DAS62, the results revealed that cultivar (p <0.001), irrigation level (p <0.001), and cultivation time (DAS, p <0.001) affected the overall trajectory of digital biomass (fig. S8A), with a significant correlation among cultivar, irrigation level, and cultivation time (p = 0.039). Water limitation affected all cultivars (p <0.05) except Cadenza. During the growing season, the digital biomass increased to a maximum value, exhibiting no treatment effect depending on the variety (p=0.01) (fig. S8A).

**Fig. 4.**
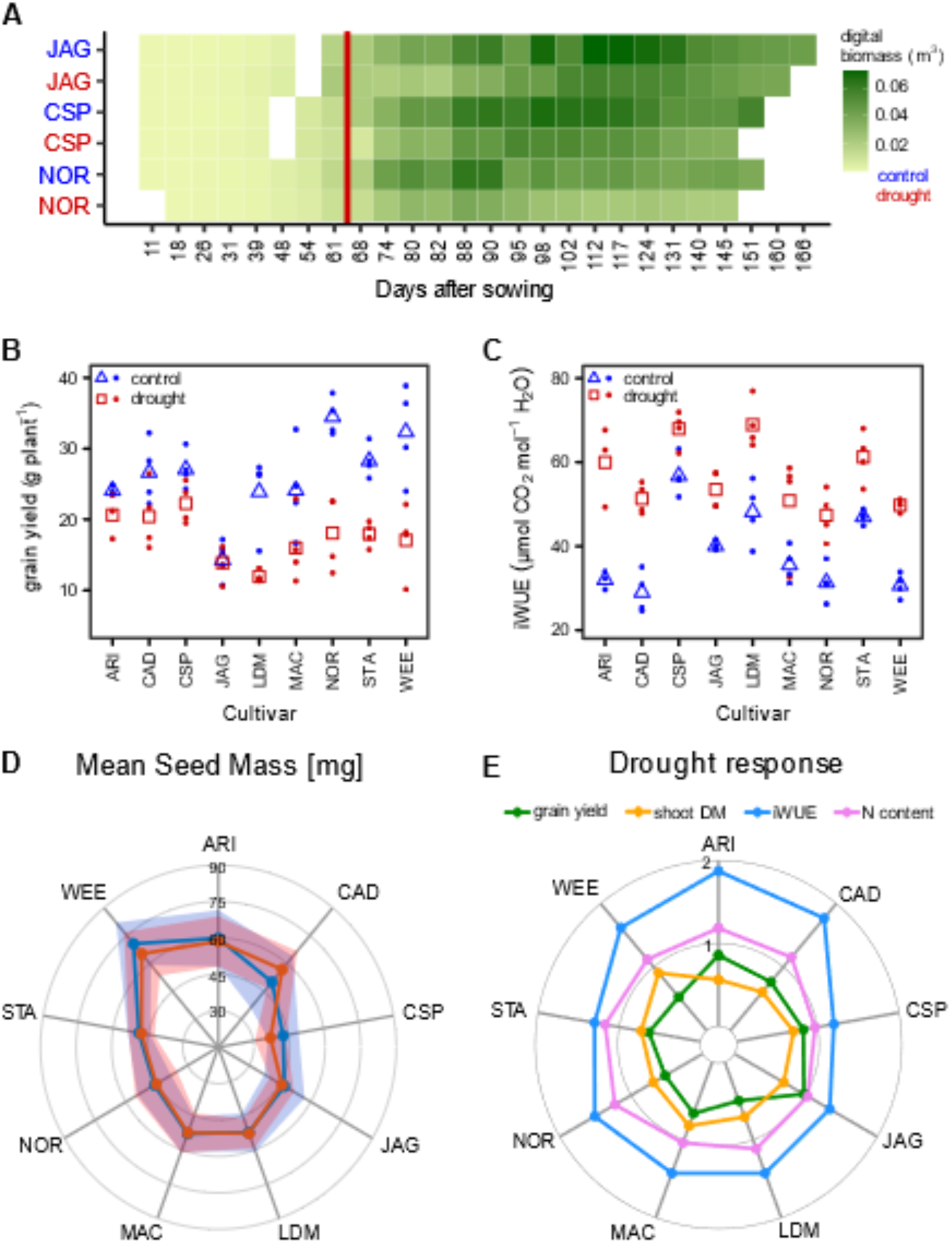
Phenotyping traits of nine cultivars from the wheat pangenome collection. (**A)** Digital biomass (m^3^) for Jagger (JAG, drought tolerant), Chinese Spring (CSP, reference cultivar) and Norin 61 (NOR, drought sensitive) throughout the experiment, water limitation started at 62 days after sowing (vertical red line). (**B**) Grain yield per plant and (**C**) Intrinsic water use efficiency (iWUE) calculated from d^13^C content of the seeds under both irrigation regimes (dots= individual plants, triangles/squares= mean values, blue= control, red= drought treatment). (**A-C**) (n=4 individual plants; Arina*LrFor* n=3). (**D**) Average mass of the seeds (mg) developed under control (blue) or drought conditions (red), shaded area represents standard deviation. (n=188 to 384, table S5). (**E**) Drought response (ratio of mean values under drought / control conditions) regarding the traits shoot dry mass (DM, yellow), grain yield per plant (green), intrinsic water use efficiency (iWUE, blue) and nitrogen content (pink) of the seeds.

The grain yield varied among cultivars (p<0.001) and treatments (p<0.001). Under conditions of sufficient water, the yield per plant ranged from 14.3 g (Jagger) to 34.5 g (Norin 61) (Fig. 4B). The drought reduced the average grain yield to varying degrees in each cultivar (Fig. 4B). While yield was almost unaffected by drought in Jagger, significant decreases were observed in CDC Stanley (p<0.001), Norin 61 (p<0.01), CDC Landmark (p<0.01), Weebill (p<0.01) and Chinese Spring (p<0.05). Correlations between grain yield and aboveground biomass were detected only for CDC Landmark (R2=0.74, p<0.01), CDC Stanley (R2=0.88, p<0.001) and Norin 61 (R2=0.81, p<0.01) (fig. S8B).

We used intrinsic water use efficiency (iWUE) as an integrative measure of physiological activity and/or acclimation to environmental conditions (Fig. 4C). The iWUE differed among cultivars (p<0.001) and increased in plants exposed to drought (p<0.001) (Fig. 4C). Under sufficient water availability, three cultivars (Chinese Spring, CDC Landmark, and CDC Stanley) had high iWUE, and five cultivars (Arina*LrFor*, Cadenza, Mace, Norin 61, and Weebill) had low iWUE; the difference between these two groups of cultivars was significant (p<0.05). Jagger presented an iWUE of 40.1 µmol CO2 mol-1 H_2_O, which was higher than that of Cadenza (p=0.023) but lower than that of Chinese Spring (p<0.001). Seed mass showed considerable plasticity, ranging from 43 mg in Chinese Spring to more than 70 mg in Weebill (Fig. 4D, fig. S9, table S5). A positive correlation (p<0.05) between irrigation and seed mass and volume was observed only for Chinese Spring and Weebill (p<0.001) and Jagger (p<0.01). In contrast, a water deficit resulted in significantly larger seeds (p<0.001) in Cadenza (Fig. 4D). Under drought conditions, iWUE (1.20-1.88), grain yield (0.50-0.97), and shoot dry mass (0.56-0.91) exhibited high phenotypic plasticity, but N content (0.97-1.23) showed low phenotypic plasticity (Fig. 4E). Weebill, CDC Landmark, CDC Stanley, Mace and Norin 61 showed a more pronounced drought response in terms of grain yield than in shoot biomass, exemplified by Weebill, which showed the least reduction in shoot dry mass (9%) despite a 47% reduction in yield.

The plant height at flowering (BBCH 61-69) varied among the cultivars (p<0.001) and was affected by drought (p<0.001). A reduction of up to 23% was observed in Arina*LrFor* and Mace (both p<0.05) and CDC Landmark, Chinese Spring, Norin 61 and Weebill (all p<0.01). Mace produced the shortest plants (average, 0.64 m under control conditions and 0.60 m under drought conditions), whereas CDC Stanley produced the tallest plants (0.88 m under control conditions and 0.84 m under drought conditions) (fig. S10).

### Grain proteome dynamics are associated with differences in reservoir material accumulation and stress-coping mechanisms following drought stress

Quantitative grain proteome measurements revealed a moderate response to drought. Differentially abundant proteins mapped to the Chinese Spring reference genome (Fig. 5A) exhibited moderate changes (adj. p value <0.05, abs. |log2FC| >1.2) in the cultivars, with the highest number of upregulated proteins under drought conditions detected in Weebill (63), Norin 61 (51) and Cadenza (44), whereas no significant increase was detected in Mace. These changes primarily affected the S-poor (CDC Stanley, Weebill and Norin 61) and S-rich (e.g., Arina*LrFor*, Cadenza, Weebill) seed storage protein accumulation (Fig. 5B), carbon homeostasis (Cadenza), negative regulation of proteolysis (Cadenza, CDC Landmark and Weebill), inhibition of protein translation through phosphorylation (Chinese Spring), reactive oxygen species (ROS) scavenging (Cadenza, CDC Landmark, Norin 61, CDC Stanley, Weebill) and osmolyte production-related protein accumulation (Norin 61, CDC Stanley, Weebill) (Fig. 5B, table S4, table S6). Although no chromosome-level enrichment was detected, in Weebill, eight proteins encoded by the storage protein loci on the short arm and long arm of chromosome 1D and five ROS scavenging-related proteins, including glutathione S-transferases (TraesCS4A02G103800, TraesCS4A02G103900), gamma-interferon-responsive lysosomal thiol protein (TraesCS4A02G112300) and Ran-binding proteins (TraesCS4A02G110400), encoded in two clusters on the short arm of chromosome 4A were upregulated. In CDC Landmark, nine proteins encoded on chromosome 5B were downregulated under drought conditions; six of these proteins were S-rich stress response proteins (ATIs, glutathione S-transferases, and xylanase inhibitors) encoded on the short arm of chromosome 5B. Other S-rich stress response proteins (ATIs, puroindoline-like protein, alkaline phosphatases (ALPs), xylanase inhibitors and endochitinases) were also downregulated, indicating an overall decrease in S-rich stress protein accumulation under drought conditions in CDC Landmark.

**Fig. 5.**
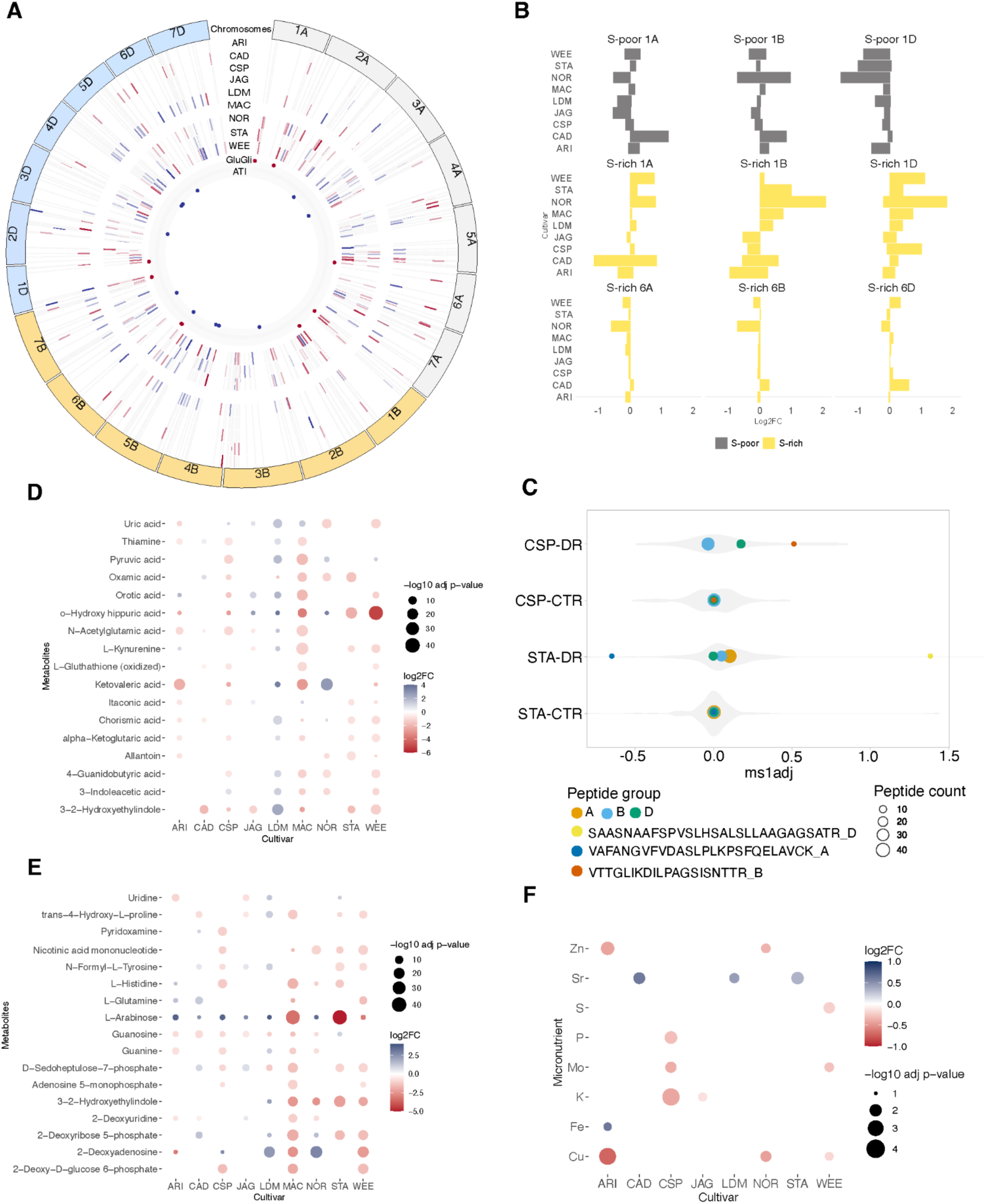
Impact of drought stress on the grain proteome, metabolome and micronutrient profiles. (**A**) Circos plot showing differentially abundant proteins identified from the cultivar-specific control vs. drought comparisons. The red line indicates high while the blue line represents low Log2FC levels due to drought. Each protein was mapped to the reference chromosomes of Chinese Spring. (**B**) Impact of drought on S-poor and S-rich gluten protein levels. (**C**) Peptide abundance biases in chromosome 5 serpin proteoforms showing upregulated peptides in subgenome D and A proteoforms in Chinese Spring during drought stress and downregulation of subgenome B peptide variant in CDC Stanley in drought conditions. (**D-F**) The dot plots show the ratio between drought and control samples for significantly different ROS scavengers (**D**), osmolytes (**E**) and micronutrients (**F**) detected from the nine cultivars.

Targeted analysis of the monitored gluten protein peptides revealed that 45% of the peptides exhibited significant variation due to drought stress and that the cultivar- and subgenome-specific drought responses in terms of the abundance of peptides representing the same gluten protein type were moderate (table S7). The cultivars Norin 61, Cadenza and Weebill showed the highest number of gluten DAPeps (23, 19 and 18, respectively), whereas no significant effect was detected in Jagger and CDC Landmark. The abundance of S-rich and S-poor gluten proteins under drought conditions differed mainly when the proteins originated from the B subgenome (Fig. 5B, table S7). In Cadenza, drought led to a significant increase in S-rich LMW glutenin accumulation (Chr 1A, 1B) but had no significant effect on the S-poor HMW glutenin and omega gliadin accumulation; however, similar to our global quantitative analysis, both S-rich and S-poor storage proteins were upregulated in Weebill (table S7). The results revealed that the effects of moderate drought manifested less robust changes in seed storage protein accumulation, but the trends indicated an unbalanced subtype accumulation, possibly due to differences in nutrient and storage protein accumulation mechanisms in response to drought stress.

The abundance of ATIs (total and by subtype) in the grains was not different between the control and drought samples for all the cultivars. However, the targeted quantitation assay identified 31 (52.54%) DAPeps between the control and drought samples (adj. p value <0.05, |log2FC| >1), primarily representing the CM3 (2 peptides) and dimeric ATI subtypes (12 peptides). Similarly, the data-independent acquisition (DIA) analysis identified 16 differentially abundant ATI proteins (adj. p value <0.05, |log2FC| >1.2), mostly representing the CM ATI subtypes (table S6). The TLR4 activation bioassay results, compared with those of the control grain samples, shows that T-cell responses were decreased in all the drought samples, but the difference was significant only for the cultivar Mace (unpaired t-test p<0.009) (table S9). ATIs are well-known for their role in the biotic stress response (*27*), while Bowman-Birk and chymotrypsin inhibitors have been reported for their activity in drought tolerance (*28*). Thus, the lower ATI quantity in drought-stressed grains suggests their minimal role in water stress conditions, as the protein degradation process may help to maintain cellular homeostasis under stress conditions, preventing toxic protein accumulation, recycling nitrogen sources, and producing amino acids for new protein synthesis (*29*, *30*). The modest decrease in TLR4 activation indicates that moderate vegetative drought stress has no major impact on the levels of immune-reactive ATIs.

#### Assessing drought responses at subgenome variation levels

Mapping the 17,253 quantified peptides to the pan-proteome resulted in 48,779 protein hits across the three subgenomes of the nine cultivars, of which 1,873 proteins (3.8%) showed significant variation under water-limited conditions. A total of 8,771 peptides detected in all three homeologous triad-derived grain proteins of any group of three cultivars were used for quantitative analysis to detect sequence variants with biased stress-specific abundance profiles. Of the 8,771 quantified triad peptides, 230 triad peptides (2.6%) showed significant differences (adj. p value < 0.1) in abundance in response to drought. These peptides mainly represented proteins involved in stress response (serpins, oxidoreductases), carbohydrate metabolism, ABA response and storage protein biosynthesis (Fig. 5C, fig. S11A, C, table S10). To analyze the overall impact of stress at the triad level, we considered proteins with an abundance that changed by at least 50%. Triad plots indicate the lack of significant bias among the three subgenomes in any of the cultivars (fig. S11B).

Serpin Z2 protein isoforms, members of the serine protease inhibitor family from the homoeologous chromosome group 5 (TraesCS5A02G417800, TraesCS5B02G419900, TraesCS5D02G425800 orthologs), that exhibited both biases in subgenome-derived proteoform use and significantly different abundance patterns due to drought stress were detected (Fig. 5C). At the triad-derived protein level, serpin Z2 was upregulated in Cadenza, CDC Landmark, Norin 61 and Weebill. The majority (92%) of the identified chr 5 group serpin proteoform peptides were shared among the subgenomes, indicating a high degree of protein conservation. The abundance of peptides with subgenome sequence variation significantly differed between the control and drought conditions in Chinese Spring (Fig. 5C) and CDC Stanley. Similarly, DAPeps were detected from 1S globulins, LEA proteins and oil body proteins (table S10, fig. S11C). Of these LEA proteins (TraesCS1A02G372700, TraesCS1B02G392700, TraesCS1D02G379300 orthologs) primarily fulfill functions related to ROS scavenging and osmolyte accumulation.

#### Impact of vegetative stage drought stress on grain small molecule and micronutrient compositions in grain

Pathway enrichment analysis of the 98 significantly up- and down-regulated central carbon pathway metabolites (adj. p<0.05; abs. |log2FC| >1) indicated that the metabolite responses primarily control photosynthesis and stomatal closure, resulting in drought avoidance, tolerance-coupled with antioxidant activities via the production of ROS scavengers and osmolyte production (Fig. 5D, 5E). These results suggest the potential role of plant stress memory in structural and physiological adaptations under water deficit conditions (*31*). The total osmolyte level significantly differed between control and stress conditions in Jagger, Mace, CDC Landmark, and Weebill. Likewise, the total abundances of ROS scavenging-related metabolites in seeds significantly differed between the conditions in Cadenza, Mace, Jagger, CDC Landmark, CDC Stanley, and Weebill. We detected significant differences in the drought response of the cultivars to osmolytes (p<0.008) and ROS scavengers (p<0.005) (Fig. 5D, 5E). For example, the ROS scavenger orotic acid synthesized through the nicotinamide-dependent redox reactions by dihydroorotate dehydrogenase (DHODH) enzyme (*32*). The overexpression of DHODH leads to increased production of orotic acid, which is involved in NO signal transduction and increases tolerance to abiotic stress conditions such as drought (*33, 34*). The increased abundance of arabinose exemplifies its role in constitutive leaf desiccation tolerance during water deficit conditions (*35*). Notably, the arabinose and galactose residues are part of the pectin-associated arabinose polymers that increase leaf cell wall flexibility upon dehydration (*36*).

The above results suggest that the role of these molecules in osmotic adjustment is to protect the cells (*37*) and stop damage at the cellular level by increasing ROS scavenging and controlling stomatal activity (*38*, *39*). Furthermore, compared with those of Mace, the cultivar CDC Landmark exhibits an increase of ∼87% in osmolyte levels and ∼116% in ROS scavenger levels, indicating that these metabolite changes improve drought adaptation and tolerance in the seeds of different cultivars (*40*). As mineral content has been reported to be affected by yield dilution (*41*), we normalised the seed micronutrient concentration by seed weight for all measured micronutrients. Comparative analysis between control and drought samples showed significant differences in key micronutrients, such as Zn (p<0.03; ArinaLrFor), K (p<0.006; Chinese Spring), Ca (p<0.03; Jagger) and P (p<0.03; Chinese Spring) accumulation increased due to drought stress (Fig 5F), aligned with the previous findings about their role in drought adaptation through ROS scavenging (*42*) or osmotic adjustment (*43*).

#### Impact of drought stress on grain protein and metabolite profiles associated with phenotypic traits and micronutrient levels

To understand how applied drought stress affects grain phenotypes in terms of proteomic, metabolomic and micronutrient compositions, network analysis was used to relate protein and metabolite modules to the measured phenotypic traits (Fig. 6, fig S12). Phenotypic traits collected during the plant growth period (0 to 169 DAS) that significantly differed (127 traits, fig. S12) between the control and drought samples were used for the trait-module association analysis, which identified significant (adj. p value <0.05) associations of protein modules with the C/N ratio, protein content, seed mass and seed length. In addition, some optical and morphological traits (e.g., DAS82 max plant height, DAS117 leaf area, DAS117 normalized pigment chlorophyll ratio index [NPCI], DAS131 leaf angle and DAS131 normalized difference vegetation index [NDVI]) (fig. S12A) were significantly related to the protein modules. Key features with significant importance identified via the variable importance in projection (VIP) values from partial least squares discriminant analysis (PLS‒DA) highlight the concerted associations of certain grain proteins with flowering time and early seed filling-related phenotypic traits. Leaf angle and area measurements at the flowering and seed filling stages (DAS102 to DAS 131) revealed strong negative correlations (r = -0.65 to -0.71, adj. p-value < 1.0e-6) with proteins involved in lipid transport, response to oxidative stress and proteolysis (fig. S12B). While proteins with positive correlations (r 0.58 to 0.66, adj. p-value <9.0e-4) are involved in storage protein accumulation, sulfur-rich enzyme inhibitor activity and protein glycosylation.

**Fig. 6.**
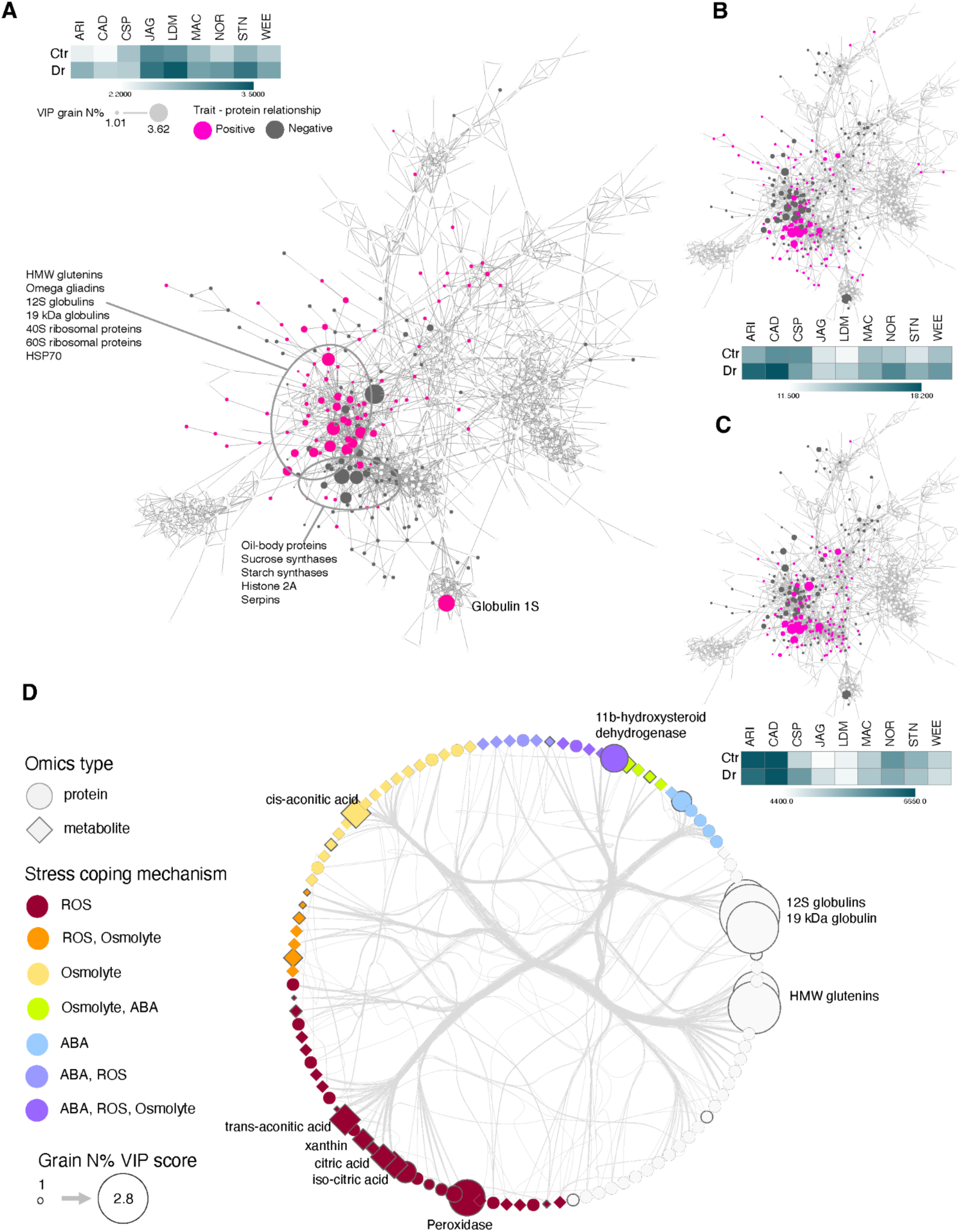
Protein co-abundance network analysis shows that osmolytes, ROS scavengers, and ABA-related proteins affect storage protein accumulation. (**A, B and C**) Protein co-abundance network highlights proteins with significant impact on selected traits of grain N% (**A**), carbon-to-nitrogen ratio (**B**) and K content (**C**). Heatmaps show the changes in the measured traits in the control and drought samples. Proteins with a positive impact on the trait are highlighted in magenta, while negative proteins are shown in grey. Dot size is proportional to the variable importance in projection (VIP) values of the selected trait. (**D**) Integrated network of proteins and metabolites overlayed with their potential function as ROS scavengers (ROS), osmolyte, ABA or a combination of these. Feature sizes are proportional to the VIP score related to grain N content.

The medium-level positive correlation between plant height (fig. S10) and a subset of grain proteins is associated with an increased accumulation of proteins with nutrient reservoir functions, oxidoreductase and other S-rich stress-related protein activities and lipid droplet organization in the seed and the accumulation of the osmolyte synthesis-related betaine aldehyde dehydrogenase (fig. S12B).

The expression of MOTHER of FT (MFT) and TFL1 protein (TraesCS7A02G229400, TraesCS7B02G195200), which is a member of the phosphatidylethanolamine-binding protein (PEBP) family with dual roles in flowering, seed dormancy and germination (*44, 45*), was increased under drought conditions (FC= 1.44, adj. p-value <0.05) in Cadenza. This protein is also a known regulator of seed oil and protein content accumulation (*46*) and is significantly correlated with grain protein content (VIP = 1.43, adj. p<0.05) and negatively correlated with K content (VIP=1.36, adj. p<0.05).

The protein serpin Z2 encoded on chromosome 5B (homeologous pan proteoforms of TraesCS5B02G419900) was significantly negatively correlated with protein content (VIP = - 7.6, adj p-value = 1.18e-5) and positively correlated with the C/N ratio (VIP = 7.39, adj. p-value = 2.97e-5) and dry weight of mature ears (VIP = 7.75, adj. p value = 3.5e-3), which reflects subgenome expression bias when drought stress is applied (Fig. 5C, table S11).

Wheat grain proteins are enriched in S-rich and S-poor storage proteins. These proteins significantly varied at the cultivar and stress levels (Fig. 2, Fig. 5B, fig. S3 and table S7). We constructed a co-abundance network to unravel the contributions of storage protein subtype variations to the measured traits (Fig. 6A-C). We found that a protein hub mostly composed of S-poor seed storage proteins (S-poor gluten proteins and 12S seed storage globulins) influenced the total protein content through an increase in S-poor storage proteins (Fig. 6A, fig. S13B, S14C), with contrasting effects on the seed C/N ratio and K content (Fig. 6B, 6C), suggesting a role for potassium in soluble sugar accumulation in the seeds. The cultivars Arina*LrFor*, Cadenza, Chinese Spring and Norin 61 accumulated high levels of carbohydrates and K under drought conditions, which maintain turgor pressure and cellular integrity to protect plants from stress (*47*) under abiotic stress conditions. Proteins with a significant positive association with K content and C/N ratio and a negative association with the grain N content (Fig. 6A, fig. S13C) primarily represent lipid content-related oleosins (e.g. TraesCS7A02G234100, TraesCS7B02G132500, TraesCS7D02G234400) and carbohydrate metabolism proteins such as sucrose synthase (TraesCS4A02G446700), further highlighting the coordinated changes in the abundance of carbohydrates and osmolyte-type micronutrients involved in drought stress.

We further examined the protein network to confirm differences in the S-poor and S-rich levels and their associated role in total protein accumulation and stress responses in the grain (Fig. 6A, fig. S13). Positive correlations (Pearson r >0.8) were detected between the S-rich gluten protein content and proteins like wheat subtilisin chymotrypsin inhibitors (WSCIs), gamma gliadins, LMW glutenins, alpha-amylase subtilisin inhibitors, xylanase inhibitors and glutathione transferases mostly involved in late seed development, biotic and abiotic stress responses and ROS scavenging (*30*) (fig. S13A). The S-rich protein hub also included the TaMFT homologs from chromosome group 3, which are related to pre-harvest sprouting (*48*). Among these proteins, an alpha-amylase subtilisin inhibitor proteins (TraesCS2B02G392500) had the strongest association with the total S-rich gluten protein content (VIP = 3.84, adj. p-value <0.05, table S11). In contrast, negative correlations (Pearson r<-0.8) were observed for S-poor proteins such as HMW glutenins, omega gliadins and 11S seed storage globulins, which were positively associated with the total grain N content (fig. S13A, 13B, table S11). Differences in K content, the C/N ratio and compositional changes between S-rich and S-poor proteins highlights the interconnectedness and co-occurrence of underlying osmolyte accumulation and ROS scavenging events in cultivars and suggest the plasticity of stress coping mechanisms under drought conditions.

Our integrated proteomic and metabolomic data (Fig. 6D) also illustrate the intricate relationships among ROS scavenging (e.g., LEA, malate dehydrogenase, citric acid, trans-aconitic acid and serpins), osmolyte production (cis-aconitic acid, xylitol and beta-hydroxysteroid dehydrogenase) and ABA regulation-related (e.g., LEA proteins, ribonuclease TUDOR1-like) proteins and metabolites in defining seed traits such as protein content, storage protein composition, or the C/N ratio. Drought stress alters protein and metabolite profiles, leading to different accumulation patterns for S-rich and S-poor proteins in the seeds, with S-rich protein abundance changes showing trends that are similar to those of ROS scavengers. In addition, the network interactions highlight the potential involvement of S-rich storage proteins and enzyme inhibitors as stress response proteins and potential antioxidants. The cultivar-specific drought responses clearly highlight differences in the stress coping mechanisms. However, cultivars such as Norin 61, CDC Stanley and Weebill accumulate increased S-poor storage proteins in seeds, which are affected by increased levels in ROS scavenging-related proteins and metabolites (fig. S14). In Arina*LrFor*, Chinese Spring and Jagger, the increased ROS scavenging activity (xanthin, uric acid and trans-aconitic acid) is not coupled with increased S-poor protein accumulation. In Chinese Spring, increased phosphorylation leads to inhibited seed protein accumulation, indicating the co-activation of protein kinases and ABA under stress to balance metabolic and cellular redox status resulting in reduced protein accumulation (*49*). Trans-aconitic acid, an abundant ROS scavenger in grasses, is upregulated in Chinese Spring, Jagger, CDC Stanley and Weebill and has a known negative impact on photosynthesis (*50*). Interestingly, CDC Landmark uses a strikingly different strategy with significantly lower levels of ROS scavenger metabolites in grains, such as trans-aconitic acid and S-rich stress proteins (ATIs, ALPs). Finally, we found that seed volume and mass were affected by the interplay of changes in abundance patterns between ROS-scavenging and osmolyte-related metabolites (Fig. 6D), which helps to maintain plant physiological processes and yield-related traits such as seed mass and volume. For example, the level of hypoxanthine, a ROS scavenger, significantly increased under drought conditions in Chinese Spring (|log2FC|=1.59, adj. p value <0.05) and Mace (|log2FC|=3.64, adj. p value <0.05). Hypoxanthine plays a role in purine and energy metabolism, is positively correlated with the osmolyte glucose and ribitol dehydrogenase (TraesCS1A02G0838001, TraesCS1B02G101100) and is positively associated with seed weight (VIP=1.39, adj. p value <0.05).

## Discussion

This study presents the first comprehensive map of grain proteins and metabolites of the wheat pan-genome cultivars, providing insights into subgenome-specific variation in protein composition and metabolite content in wheat grains and the ability to adapt to soil water deficiency. Phenotypic measurements revealed the plasticity at the cultivar level in terms of shoot biomass and height, iWUE, grain yield and grain N content, which define the cultivar-specific response to drought stress (Fig. 4). Our comparative peptide level analysis demonstrated that the core grain peptidome accounts for 50% of the matching peptides, shared by all cultivars. Notably, the shared and unique peptides that play prominent roles in modulating key biological functions such as protein synthesis, folding and accumulation and the stress response, explain the observed variation at the cultivar level. In contrast, the core peptidome is associated with primary metabolic processes. This finding is consistent with those of previous pan-genome studies in wheat (*9*), *Arabidopsis thaliana* (*51*) and soybean (*52*). The results of this study indicate balanced expression of homeologous proteins in the A, B and D subgenomes, with peptide specificity at the subgenome level associated with the cultivar phenotypes (Fig. 1 and Fig. 5). The protein content and composition patterns of the grains revealed differences in the S-rich and S-poor seed storage protein accumulation patterns associated with the vegetative and flowering stage phenotypes (fig. S12). Plant height was positively correlated with protein content, especially with S-poor protein levels, suggesting that their variation is controlled by cultivar genetics, including phenology genes such as MFT, which has dual roles in flowering and late seed development and germination (*53*).

We have developed a pan-wheat multiome resource that integrates information at multiple levels and can aid in the identification of genes, small molecules and pathways associated with measured traits under both control and moderate drought stress. Our high-resolution proteomic analysis provides a new understanding of how polyploidization and cultivar differences affect phenotypic and biochemical traits and wheat grain composition under moderate drought stress. In particular, the described cultivar differences in the subgenomes can aid in the selection of suitable cultivars for different climatic conditions (*54*) or improve selection for tailored breeding purposes (*55*).

The drought adaptation and tolerance mechanisms identified are consistent with previous findings on changes in iWUE, transpiration rate and phenology (*56*). Plants protect themselves against water shortages through a coordinated interplay of metabolites, proteins and micronutrients by adjusting their cellular osmotic potential and accumulating biomolecules such as sugars and amino acids to maintain water retention, provide membrane protection, and act as antioxidants against ROS in different cell compartments (*57*). ABA is primarily involved in closing the stomata to reduce the transpiration rate but it is also interlinked with branched-chain amino acids and sugars to control ROS production and sugar-sensitive signalling under drought stress (*58*), which is evident in our network analysis (Fig. 6). In an exemplary set of cultivars, Arina*LrFor* and Chinese Spring, we detected cultivar-specific differences in the accumulation patterns of osmolytes, ROS scavengers and photorespiration-associated metabolites for drought tolerance (Fig. 6D). The drought-responsive wheat cultivars (Arina*LrFor*, Cadenza, CDC Landmark and CDC Stanley) showed high osmotic and stomatal adaptations (*59*) but low response to ABA (*60*) (Fig. 6). Furthermore, the cultivars Chinese Spring, Cadenza and Norin 61 showed moderate drought tolerance, which is reflected in their minimal osmotic adjustment potential (Fig. 5D), resulting in increased water-soluble carbohydrates that can be remobilized for grain filling and yield maintenance under drought conditions. Interestingly, the opposite trend between the accumulation of ROS scavengers and the osmolytes was observed when we compared the metabolites and networks (Fig. 5E), as shown in the cultivars Arina*LrFor* and Chinese Spring. These results suggest that Chinese Spring seeds accumulate osmolytes and ABA-linked metabolites to maintain turgor pressure (*61*), whereas Arina*LrFor* uses the ROS-scavenging metabolites in combination with glutathione-like proteins to neutralize free radicals (*62*), further indicating the drought response plasticity of these cultivars.

The adaptable stress-induced shifts in omics-level data in mature grains demonstrate the importance of integrating crop genetics, environmental factors and multilevel omics measurements in selecting suitable crop varieties (*11*). While our study provides valuable insights, further research with larger cultivar sets grown under different climatic conditions, analyzing additional plant tissues such as roots and leaves, post-translational modifications, and phenological and quality trait measurements, would more comprehensively address the complexity of drought adaptation mechanisms, as implemented in recent studies (*63*).

In conclusion, our work shows that subgenome-specific peptide abundance patterns in polyploid bread wheat grains are associated with stress-responsive variation. These patterns indicate cultivars’ ability to employ stress-coping mechanisms through cellular divergence and the concerted interplay between biomolecules to determine functionality and plasticity at the subspecies level. These comprehensive datasets and analyses provide a framework to help researchers and breeders develop strategies to improve crop resilience by manipulating homologs to modulate trait response.

## Materials and methods

### Origin of plant material for multi-omics analyses

The seeds for the multi-omics analyses were obtained from an experiment on the high-throughput phenotyping platform (HTP) (‘FitnessSCREEN’) in the greenhouse of the Environmental Simulation Research Unit (EUS) at Helmholtz Center Munich in Neuherberg, Germany (fig. S5). The experiment (see also section below) was conducted on nine varieties of wheat (*Triticum aestivum* L.) derived from the genome assemblies of the 10+ Wheat Genome Project (*6*).

### Plant growing conditions and phenotyping of morphological traits

The plants were grown from seed to maturity in a ’pot-in-pot’ system consisting of a small pot (0.8 L) with bottom openings placed in a large pot (8.5L), one plant per pot (fig. S5). This ’pot-in-pot’ system initially limited the rooting space of the seedlings according to the plant density and competition situation in the field but provided enough space for roots to expand during growth, thus minimising ’pot binding’ (*64*). The pots were filled with a mixture of 45.5% peat substrate (Einheitserde CLT, Balster Einheitserdewerk, Fröndenberg, Germany; N 250 mg L^-1^, P_2_O_5_ 300 mg L^-1^, K_2_O 400 mg L^-1^) and 54.5% sand (0.6 - 1.2 mm size). Prior to the experiment, the maximum water holding capacity of the pots (’pot capacity’) was determined (*65*). From 77 days after sowing (DAS), 0.5% of a N:P:K (3:27:18) fertilizer (“Phosphik" Biolchim, Hannover, Germany) was added to the irrigation water once a week.

The climatic conditions in the greenhouse followed the outdoor conditions throughout the experiment from the beginning of March to end of July 2020 (fig. S6). The daylight duration was initially set to 12 h, but followed the outdoor conditions once the photoperiod exceeded 12 h per day (fig. S6A). To ensure optimal growth, we supplemented the natural solar radiation in the greenhouse with a combination of HP sodium vapour lamps (SOD Agro 400, DH Licht GmbH, Wülfrath, Germany) and quartz metal halide lamps (MASTER HPI-T Plus 400W, Philips, Eindhoven, The Netherlands) when the radiation intensity outside the greenhouse fell below 20 klx (corresponding to 370 µmol m^-2^ s^-1^ photosynthetic photon flux density (PPFD)) during the day (7 am to 7 pm) (fig. S6B).

At the beginning of the experiment, all plants of each cultivar were grown under the same water regime (fig. S7A). At DAS 61, when most of the plants had reached BBCH stage 32, half of the plants were kept under well-watered conditions (’control’, n=4), while the other half was subjected to moderate drought stress (’drought’, n=4). Drought was induced by withholding irrigation until DAS 77. Thereafter, plants received the amount of (deionized) water needed to reach the desired target weight every 3 to 4 days (fig. S7A), which corresponded to 75% of pot capacity in the control treatment and 40% of pot capacity in the drought treatment.

Agronomic traits were assessed once or twice a week on the HTP platform, where up to 5 pots were placed in a carrier (fig. S5). Carriers were automatically transported to the measurement station. Plants were individually scanned by a multispectral 3D dual scanner (PlantEye F500, Phenospex, Heerlen, Netherlands) mounted 1.35 m above the plot edge and moved over the plants (fig. S5D). The PlantEye produced a 3D point cloud from which 3D plant models were created using the x, y and z coordinates (voxels) of each point (fig. S5F). From the 3D models, all plant parameters such as plant height, leaf area, digital biomass (height x leaf area) were calculated and stored in the HortControl software (Phenospex, Heerlen, Netherlands). To eliminate possible positional effects in the greenhouse, the carriers were repositioned in the greenhouse cabin after each scan. At the beginning of the drought treatment, the number of pots per tray was reduced to three to avoid overlapping of leaves and to allow good separation of individual plants.

Plants were harvested at maturity (BBCH ≥ 92). Morphological parameters of the above-ground parts (maximum plant height, number and length of spikes, shoot dry mass) were determined manually. Spikes were discarded and the total grain yield per plant was determined (Fig. 4B). Thousand-grain weight were determined on a set of 100 grains for each plant (fig. S9C).

Individual seed characteristics (two-dimensional projected area [mm²], length [mm], width [mm], mass [mg] from weighing (fig. S9), volume [mm³] from volume carving (*66*) and density from the ratio of seed mass to volume [mg/mm³]) were evaluated in the phenoSeeder (*67*) at Forschungszentrum Jülich Jülich, Germany. Seed material from plants (n=4, except Arina*Lr*For n=3) of each genotype and treatment level was analyzed. The number of individual seeds for which all traits could be successfully determined ranged from a minimum of 188 for Landmark to a maximum of 383 for Norin 61.

Carbon (C) and nitrogen (N) content, as well as the natural abundance of the stable isotopes ^13^C and ^15^N in the grain, were determined in 1.3 mg aliquots of 10 homogenized grains per plant using an isotope ratio mass spectrometer (IRMS) (delta V Advantage, ThermoFischer, Dreieich, Germany) coupled to an elemental analyzer (EURO EA, Eurovector, Milan, Italy) (*68*). A lab standard (acetanilide), included in each sequence at intervals and different weights, was used to calibrate and to determine the isotopic linearity of the system. The laboratory standard itself was calibrated against several suitable international isotopic standards (IAEA; Vienna).

Grain protein content was calculated from total N content using the conversion coefficient of 5.7 (for the wheat endosperm) (https://archive.org/details/factorsforconver183jone), see also (*69*).

#### Determination of Water Use Efficiency (WUE)

Based on the carbon isotope signature d^13^C of the grain and in the atmosphere (approximately -8 ‰), the carbon isotope discrimination (D^13^C) was assessed according to (*70–72*). From these values and the atmospheric [CO_2_] in the greenhouse during the experiment (average 420 µmol mol^-1^ CO_2_), the intrinsic water use efficiency (iWUE) was calculated, which provides integrative information not only on the environmental conditions but also on the physiological activity during the entire vegetation period (*73*, *74*).

### Proteomics sample preparation for wheat samples

#### Protein extraction, estimation, and digestion for Data Dependent Acquisition (DDA) and Data Independent Acquisition (DIA)

The wheat flour (10 mg) from nine cultivars was weighed into 1.5 mL low protein bind tubes (Eppendorf, Hamburg, Germany) in four replicates. One hundred µL (10 µL/mg) of protein extraction solvent consisting of 8 M urea/2M thiourea/4% CHAPS/50 mM DTT/0.1 M Tris-HCl at pH 8.4 was added with vortex mixing until the flour was thoroughly mixed with the solvent. The tubes were then sonicated for 5 min and incubated on a thermomixer (Eppendorf, Hamburg, Germany) at 600 rpm for 60 min at 22°C. The tubes were centrifuged for 10 min at 20,800 ×g, and the supernatant was transferred to a fresh tube (Eppendorf, Hamburg, Germany). The protein concentration was estimated using a Bradford colorimetric assay (Bio-Rad, South Granville, Australia), following the manufacturer’s protocol, to calculate the volume equivalent of 100 µg of protein before loading onto a 3 kDa molecular weight cutoff (MWCO) filter (Merck Millipore, Bayswater, Australia). Tryptic peptides were prepared using the FASP protocol as previously described (*75*). In short, the protein on the filter was washed twice with 8 M urea in 0.1 M Tris-HCl buffer (pH 8.5) and centrifuged for 15 minutes at 20,800 × g. For cysteine alkylation, 50 mM of iodoacetamide (dissolved in 8 M urea and 100 mM Tris-HCl) was added to the filters. The mixture was then incubated in the dark at 22°C on an Eppendorf ThermoMixer® C (Eppendorf, Hamburg, Germany) at 600 rpm for 30 minutes followed by centrifugation at 20,800 x g for 10 minutes. The buffer was exchanged with 100 mM ammonium bicarbonate (pH 8.0) by two consecutive wash/centrifugation steps. The sequencing grade digestion enzyme, trypsin (Promega, Alexandria, Australia) solution (2 μg in 200 μL, 10 μg/mL in 50 mM ammonium bicarbonate and 1 mM CaCl_2_) was applied to the filter and incubated for 18 h at 37°C in a thermomixer at 300 rpm. The tryptic peptides were collected by centrifugation at 20,800 x g for 15 minutes. They were then washed twice with 200 µL of 50 mM ammonium bicarbonate and the combined filtrates were lyophilized. The digested peptides were stored at -20°C until resuspension in 100 μL of 1% formic acid containing 0.1 pmol/µL indexed retention time (iRT) peptides for analysis. Two pooled biological quality control (PBQC) samples were prepared prior to storing by aliquoting 2 µL of reconstituted digested peptides from combining all samples and cultivars with their respective treatment groups. The combined PBQC sample was injected first using DDA for DIA window calculation and later to evaluate the data acquisition quality during the DIA experiment. The sample and treatment-specific PBQCs were used for the DDA experiment.

#### Protein extraction and estimation for glutenins, gliadins and ATIs for MRM-MS

Ten mg of flour samples in four replicates from nine wheat cultivars were measured into 1.5 mL low bind microtubes (Eppendorf, Hamburg, Germany). The extraction buffer consisted of 55% (v/v) propane-2-ol (IPA) and 2% (w/v) DTT in water. The protein extraction solvent was added to each tube and the flour samples were vortexed, followed by sonication for 5 minutes at room temperature until thoroughly mixed with the flour. The sample tubes were incubated for 30 minutes at 400 rpm at 50°C. The extraction solution containing tubes were then centrifuged for 15 minutes at 20,800 *g*. Protein concentration was estimated by Bradford assay and 50 µg protein was transferred to 3 kDa MWCO filters for on-filter protein reduction, alkylation and digestion with trypsin and chymotrypsin (50:1 protein:enzyme ratio) (Sigma-Aldrich, Bayswater, Australia). The detailed protein extraction and sample clean-up steps have been described previously (*76*, *77*). Two separate PBQC samples were prepared from trypsin and chymotrypsin digested samples for glutenin and gliadins, and ATIs by aliquoting 2 µL of sample from each replicate. These pooled samples were primarily used to refine the MRM method.

#### ATI extraction and bioactivity testing on HeLa TLR4 dual reporter cells

ATIs were quantitatively solubilized from 100 mg of wheat flours in 2 consecutive extractions with 0.5 ml of 10 mM Tris, 0.5M NaCl, pH 7.8, both supernatants were combined, and 10 µl were tested for TLR4 stimulating bioactivities using aTLR4/MD-2/CD14/IL-8 Prom/LUCPorter™ reporter cell line (Novusbio, Wiesbaden, Germany), that expresses Renilla luciferase signals under the control of an IL-8 promotor. The cell line was further transfected with a pCMB-firefly-luc-hygro expressing firefly luciferase to measure cell viability as previously described (*78*). Prior incubation of extracts with polymyxin B ruled out LPS (lipopolysaccharide, a strong activator of TLR4) in all extracts. To calculate the normalized TLR4 activity, the TLR4-driven Renilla luciferase signal was divided by the constitutive Firefly luciferase signal RLU, relative luminescence units) yielding normalized TLR4 stimulating bioactivities which were further normalized to a wheat flour extract standard curve based on 3 mg of bioactive ATI protein per g of the dried standard wheat flour.

#### Database construction

Translated gene models of the pan-wheat cultivars (*6*) were appended with Biognosys iRT pseudo-protein sequence and sequences representing the common repository of adventitious proteins (cRAP). Homologous sequences were clustered using CD-HIT v.4.7 with a sequence identity threshold of 0.95 (*79*). The final search database contained 567,994 protein sequences.

#### Data-dependent acquisition

The digested wheat samples were reconstituted in 100 μL of 1% formic acid. Cultivar- and treatment-specific PBQC samples were prepared from the four replicates of each wheat variety for proteome measurements using gas-phase fractionation-data dependant acquisition (GPF-DDA). In brief, 2 µL of the sample was injected into an Ekspert nanoLC 415 (Eksigent, Dublin, CA, USA) operating in a trap-elute configuration. The mass spectrometry data was acquired across two *m/z* windows: 350-650 and 650-1250 over a 55-minute LC gradient. The injected samples were chromatographically separated on an Ekspert nanoLC415 (Eksigent, Dublin, CA, USA) directly coupled to the OptiFlow ion source of a TripleTOF 6600 LC-MS/MS (AB SCIEX, Redwood City, CA, USA) system, as previously described (*76, 77*). In short, the digested peptide samples were desalted for 5 minutes using a ChromXP C18 (3 μm, 120 Å, 10 × 0.3 mm) trap column at a flow rate of 10 μL/min with solvent A, and then separated on a ChromXP C18 (3 μm, 120 Å, 150 mm × 0.3 mm) column at a flow rate of 5 μL/min. The solvents were: (A) 5% DMSO, 0.1% formic acid, 94.9% water and (B) 5% DMSO, 0.1% formic acid, 90% acetonitrile and 4.9% water. A linear gradient from 5% to 45% solvent B over 40 minutes was employed, followed by 45% to 90% B over 5 min, a 5-minute hold at 90% B, returning to 5% B over 1 min, and 14 min of re-equilibration. The ion spray voltage was set to 4500 V; the curtain gas was set to 30 psi, and the ion source gases 1 and 2 (GS1 and GS2) were set to 30 psi. The heated interface was set to 150 °C.

#### Proteome measurements using data-independent acquisition

A master pooled sample was prepared by combining the cultivar-level sample pools and used to acquire a single-shot DDA experiment with a mass range spanning *m/z* 350-1250. The distribution of precursors across the overall retention time-averaged mass spectrum was then used to calculate optimum variable mass windows for DIA-MS acquisition using the sequential window acquisition of all theoretical mass spectra (SWATH-MS) Variable Window Calculator v1 (AB SCIEX, Redwood City, CA, USA). DIA experiment windows were calculated using the following parameters: *m/z* range 350-1250, accumulation time 250 ms and 100 variable windows with a 40 ms accumulation time. A cycle time of 3.1004 s gave at least nine points across a chromatographic peak at half height. To acquire the DIA-MS data, the Ekspert nanoLC 415 (Eksigent, Dublin, CA, USA) was again loaded with a 2 µL injection volume per sample, operated in trap-elute configuration, with the same columns and gradients as described for DDA, and eluted directly onto the TripleTOF 6600 (AB SCIEX, Redwood City, CA, USA) MS. To minimise batch effects due to data acquisition, we randomised the sample orders during data acquisition and a blank injection was included between samples to minimise column carry-over. The PBQC samples were included within the batch to assess the instrument performance over the data acquisition period.

### Proteomics data processing and data matrix preparation

#### Data-dependent acquisition

The discovery data files of replicate cultivar pools were searched against the created pan-wheat database using the SCIEX OneOmics suite with ProteinPilot v.5.0.3.360 software. ProteinPilot search parameters were described in detail previously (*76*). The search parameters included cysteine modified by iodoacetamide, biological modifications and trypsin as the digestion enzyme. The protein and peptide identifications were performed using a 1% global false discovery rate (FDR) cut-off, respectively, calculated by the in-built Paragon and ProGroup algorithm (*80*).

#### Data-independent acquisition data processing using DIA-NN software

DIA-NN, a neural network and interference correction software for data-independent acquisition proteome measurement, was used to process the DIA-MS data (*81*). The data was searched in a library-free mode against the clustered pan genome database. Fully tryptic peptides of a length of 7 to 30 amino acids and one missed cleavage were used in the analyses. As a fixed modification, carbamidomethylation of cysteine was selected, and no variable modifications were allowed. Precursor *m/z* range was selected as 350-1250, fragment ion *m/z* range was 200-1800 and 1% precursor FDR was used for filtering. Mass accuracy, MS1 accuracy and scan window were selected in automatic modes, and interferences predicted by the software were removed. The neural network classifier was run in single-pass mode. High accuracy was selected as a quantification strategy, and cross-run normalisation was performed in a retention time-dependent manner.

#### Data pre-processing, missing value imputation and filtering

DIA results were processed both at protein and peptide levels. Peptides with abundance values <10 in all samples were excluded from the subsequent data analysis steps. Missing values were imputed in MetImp using the MCAR/MAR algorithm by considering cultivar-specific variations (82). Missing values consistently present in all four replicates of a cultivar were replaced by 0 and not imputed.

#### Peptide mapping and genome visualization

Distinct peptide lists filtered from the GPF-DDA and DIA analyses were mapped to cultivar-specific wheat proteomes (*6*) and the Chinese Spring reference genome IWGSC CS v1.1 (*7*) using the Motif search algorithm with 100% sequence matching in the CLC Genomics Workbench v24 (Qiagen, Aarhus, Denmark). The core, shell and cloud peptidomes were defined by identifying peptides that are characteristic at each cultivar (core), a subset of cultivars (shell) or a single cultivar (cloud peptidome) and visualised using Circa 1.0 (OMGenomics Labs, https://circa.omgenomics.com). Pre-processed DIA peptides were mapped to the cultivar-specific pan proteomes, and proteins with a minimum number of two quantified peptides were used for the statistical and pangenome comparisons. Mapped differentially abundant proteins were visualised on the IWGSC CS v1.1 reference genome using Circa 2.0 (OMGenomics Labs, https://circa.omgenomics.com).

#### Triad proteoform analyses

Translated sequences of orthologous genes for each cultivar were identified as reported in (*9*). The protein sequences representing homeolog gene triads of A, B and D subgenome copies with at least two quantified peptides per homeologous proteins were filtered for further analysis. Results were plotted in R using the packages ggplot2 v.3.5.1, ggtern v.3.5.0, dplyr v.1.1.4 and ggsci v3.2.0.

We used a strategy called peptide correlation analysis (PeCorA) to identify discordant peptides for differentially regulated proteoform changes across the subgenomes and treatment groups by mapping peptides to a single protein isoform (*83*). The PeCorA package uses a linear model to assess if a peptide’s variation across treatment groups differs from other peptides of the same protein. For the analysis, proteoforms of the same triad were labelled based on the subgenome allocation and change in each peptide was compared to all other peptides assigned to the same triad proteoform.

#### Gliadin, glutenin and ATI sequence selection for developing targeted quantitation assays

The glutenin, gliadin and ATI protein sequences were retrieved from the wheat pan-genome resources using homology search with the protein sequences identified in the IWGSC CS v1.1 reference genome (*14*). Protein sequences were exported to Skyline software (*84*) to develop the multiple reaction monitoring mass spectrometry (MRM-MS) assays. For glutenin and gliadin-like proteins, trypsin and chymotrypsin enzymes were used for *in silico* digestion, while only tryptic peptides were generated for the ATI proteins. Peptides were selected based on the highest intense peaks and were fully tryptic, with carbomidomethyl (C), oxidation (M) and pyro-glu N-term (Q) modifications allowed. The peptide selection, method refinement and final data acquisition were described previously (*20*, *77*). In short, unscheduled methods were exported from Skyline software to refine the MRM methods using a pooled sample. MRM data were acquired from a pooled sample and injected three times to evaluate reproducibility. The results obtained were used to refine the transitions and schedule retention times. The peptides were selected when, yielding intense peaks, with fully tryptic termini, selected modifications (M and Q) and no missed cleavages. Monitored peptide sequences were aligned to the protein sequences to define peptide markers for gluten protein-, ATI subtype and subgenome and chromosome-specific differentiation.

#### Peptide-level functional analysis using UniPept

Distinct tryptic peptide sets obtained from the filtered result sets of the GPF-DDA and DIA-NN analyses were used for functional overrepresentation analysis in Unipept Desktop v. 2.0.2 (85). GO Biological Process, GO Cellular Component, GO Molecular function and Enzyme Commission terms were analysed. Overrepresentation results were visualised using column-normalised heatmaps.

#### Central carbon metabolism metabolites (CCMM) quantitation

The targeted analysis of CCM metabolites was conducted via Agilent Infinity Flex II UHPLC coupled to an Agilent 6470 Triple Quadrupole Mass Spectrometer (LC-QQQ-MS) (Agilent Technologies, USA) as previously described in (*86*). In short, 20 mg of grounded wheat flour was extracted with 100 μL MilliQ Water and 450 μL of ice-cold (−20 °C) methanol: ethanol (1:1 v/v; LiChrosolv®, Merck, Darmstadt, Germany) spiked with 1 ppm Succinic Acid ^13^C_2_ (Cambridge Isotope Laboratories, Inc., Tewksbury, MA, USA). The mixtures were vortexed, then precipitated at -80C for an hour and then centrifuged (Centrifuge 5430 R, Eppendorf, Hamburg, Germany) at 14,000 g at 4 °C for 5 min. The supernatant was collected and filtered with Captiva EMR cartridges (40 mg, 1 mL; Agilent Technologies, Australia) to remove the lipid fraction using a positive pressure manifold (Agilent PPM48 Processor, Agilent Technologies, Santa Clara, CA). It was followed by a cartridge washing step with two 200 μL aliquots of MilliQ water: methanol: ethanol (2:1:1, v/v/v). The filtered supernatant and cartridge washes were collected into a 1.5 mL high recovery vial (30 μL reservoir, silanised glass vials, Agilent Technologies, Australia) and dried in a SpeedVAC (10 mBar). The metabolite fraction was reconstituted with 100 μL MilliQ water: methanol (4:1, v/v) spiked with 100 ppm of L-Phenylalanine (1–13C) and measured on an LC-QQQ-MS. The residual relative standard deviation (RSD) of the internal standards was 0.8% (Succinic Acid, 1,4–^13^C_2_) and 4.3% (L-Phenylalanine, 1–^13^C). Procedural blanks (n = 6), amino acid and organic standard mixtures, and pooled biological samples (n = 6) were randomly dispersed throughout the acquisition. The RSD of mixed amino acid and organic acid standards over the analytical sequence is presented in the parenthesis for Mevalonic acid (2.5%), L-Phenylalanine (4.4%), Succinic acid (3.3%), L-Aspartic Acid (3.3%), L-Glutamic acid (2.8%), L-Methionine (6.0%), L-Threonine (2..3%), L-Serine (3.0%), L-Cystine (2.4%), Lactic acid (4.9%), L-Arginine (2.6%), and L-Histidine (4.3%). All PBQC samples (n = 6) were within 10% RSD; blanks exhibited no sample carryover.

#### Sample preparation and data acquisition using ICP-MS for micronutrient analysis

The dried and ground wheat samples (approximately 0.5 g) were weighed directly into a 75-mL borosilicate glass digestion tube (Quickfit™, Thermo Fisher). A 5 mL concentrated HNO_3_ (trace analytical grade, 70%) was added into the tube and the tube containing the mixture was put on rest in a fume hood overnight. The next morning, the tubes were heated in a block system (BD 50, Seal Analytical) with a validated heating and cooling program (*87*). The tubes were withdrawn from the block when only a small residual liquid was seen, cooled under fume hood and diluted to a 10 mL capacity while a mild vortex was applied to harvest all residual from the glass tube. The mixture was then filtered by 0.45 µm cellulose acetate syringe filter and the filtrate was transferred into a pre-clean plastic tube for the analysis. All the elements except Fe were measured by an inductively coupled plasma mass spectrometry (ICP-MS, 7900, Agilent Technologies, Tokyo, Japan). Iron was measured by inductively coupled plasma optical emission spectrometry (ICP-OES, Avio 200, Perkin Elmer Instrument, USA). The preparation of standard calibration curves of different elements for the ICP-MS and ICP-OES have been described in a recent study (*88*).

Standard reference materials (SRMs) of rice flour (1568b) and tomato leaves (SRM 1573a) from the National Institute of Standards and Technology (NIST) were used to verify the results for elements in rice-based samples. Analytical results of trace elements in rice flour and tomato leaves indicated that the observed values were very close to the certified values.

#### Statistical analysis and data visualisation

#### Analysing and correlating targeted protein measurements and immunoassays

To calculate glutenin, gliadin and ATI subtypes across wheat cultivars and treatments, the average intensity was calculated for each subtype and compared to the fold-change for the highest and lowest prolamin subtype contents using Excel (Microsoft) and GraphPad Prism (v 9.3.1). In the present study, all values are presented as mean ± sd, and the figure legends mention the numbers (n) of samples or replicates. Significance levels of differences were calculated with GraphPad Prism (v 9.3.1) software using one-way ANOVA followed by Tukey’s test for multiple-group comparison and are indicated with *p* values, an asterisk (*) or numbers. Data management and visualisation were performed in Python using Google Colab. NumPy v1.21.6 was used for numerical computations and array manipulations. Pandas v1.3.5 was used for data preprocessing, filtering and tabular manipulations. Matplotlib v.3.4.3 was used to create plots and customize visualisation. Seaborn v0.11.2 was employed to generate advanced, statistically informative visualisations.

#### Weighted gene coexpression analysis, integrated analysis and network visualization

Filtered DIA-NN data was log 10 transformed and analyzed using the WGCNA package (*89*) in Mibiomics (*90*). Modules showing a strong significant correlation (p-value <0.05) with the assessed plant and grain phenotype results, total S-rich and S-poor gluten protein content and micronutrient composition were further analyzed. Partial least square discriminant analysis (PLS-DA) and variable importance in projection (VIP) were used to define the proteins significantly contributing to the selected modules. Unique proteins and peptide groups with adjusted p-value <0.05 and VIP values >1 were considered important variables for GO enrichment analysis and network visualization. R package ggplot2 was used for data visualisation.

Proteomics and metabolomics results were integrated using OmicsAnalystR (*91*). The missing value imputed and preprocessed proteomics and metabolomics data were Pareto scaled. Significantly different features were identified at p-value cut-off 0.05 and the multiblock PLS-DA method (DIABLO) (*92*) was used to identify key molecular drivers for network analysis. Pairwise similarity between selected features was calculated using Pearson correlation and filtered for correlations between the two omics datasets with a correlation threshold of 0.7. The resulting network was downloaded and further processed for visualisation in Cytoscape v. 3.10.2 using Group Attributes layout.

Protein co-abundance networks were created in Cytoscape v. 3.10.2. using the Expression Correlation App and undirected analysis with 0.8 Pearson correlation coefficient cut-off. Networks were visualised using the Force-directed layout. Proteins associated with selected phenotypes with a VIP score higher than one were annotated in the networks.

#### Statistical analysis of phenotypic traits

For statistical analysis of phenotypic traits we used R version 4.3.1 and the R package rstatix v. 0.7.2. Plots of phenotypic traits were generated using the R packages ggpmisc v. 0.6.0, ggpubr v. 0.6.0, ggradar v. 0.2, svglite v. 2.1.3, tidyverse v. 2.0.0 (*93*), and MATLAB (MathWorks, Aachen, Germany).

Residual analysis was performed to test the assumptions for either two-way or three-way ANOVA. Outliers were identified, normality was assessed using Shapiro-Wilk’s normality test and homogeneity of variances was assessed by Levene’s test. A three-way ANOVA was conducted to determine the effects of cultivar, irrigation level and time on digital biomass after onset of drought treatment. The effects of irrigation regime and time (in terms of DAS) on digital biomass of each cultivar were determined by a two-way ANOVA. To determine the effects of cultivar and treatment on the traits grain yield, intrinsic WUE, and plant height at anthesis a two-way ANOVA was performed which were followed by post-hoc multiple comparisons using the Tukey HSD. Welch’s t-test was performed to determine treatment effects on seed mass and volume distributions.

## Acknowledgements

The authors are grateful to access the proteomics facilities at Edith Cowan University (ECU) and Commonwealth Scientific and Industrial Research Organisation (CSIRO). We also would like to thank the team at the EUS greenhouse at the Helmholtz Zentrum München for carrying out the wheat experiment.

## Funding

This work was supported by the Federal Ministry of Education and Research, Germany in the frame of the German Network of Plant Phenotyping (DPPN, no. 031A053A and 031A053C), the Australian Research Council Centre of Excellence for Innovations in Peptide and Protein Science (CE200100012), Coeliac Australia (G1005443), Australian Research Council Discovery Project (DP210101705) and the Edith Cowan University School of Science Early and Mid-Career Research Grant Scheme 2022. The authors also thanked to CSIRO Early Research Career (CERC) postdoctoral fellowships.

## Author Contributions

CP supplied seeds for this experiment. ES, JBW & AS performed the phenotyping experiment, collected and analyzed experimental data and supplied materials for ‘omics experiment. FB prepared samples and performed elemental and IRMS analysis, JBW performed phenomics analysis, GH & RK performed seed analysis and analyzed data. UB, SEC, SS, ALD, JB & KB prepared proteomics samples, developed acquisition methods and collected proteomics data. UB, AJ and SAM analyzed and visualized the omics data. BB & MMR performed ICP-MS analysis and analyzed micronutrient data. DJB prepared metabolomics samples and collected and analyzed metabolomics data. MN and DS designed and performed the ATI bioassay experiments and reviewed the manuscript. UB, JBW, AJ, MS, GH, RK & JPS contributed to preparing the first draft. KFXM, MLC, CP & DS provided intellectual input. JPS, UB, MLC & AJ provided funding. JPS, JBW, KFXM, MS, UB & AJ conceptualized the project. All co-authors have critically revised the manuscript.

## Competing interests

The authors declare no competing interests.

## Data and materials availability

The raw and processed targeted proteomics data (MRM-MS) were deposited to Peptide Atlas part of the ProteomeXchange consortium, and can be accessed using the following link http://www.peptideatlas.org/PASS/PASS05899. The username: PASS05899 and password: DF5975gb. The raw and processed data independent acquisition and data dependent data acquisition datasets were deposited to ProteomeXchange consortium via PRIDE website (project accession number: PXD058792).

The username: and password: kzK4Nv8Xi2uu. Phenotypic trait data are available on OSF repository website by this link (https://osf.io/pd3ec/?view_only=eb96f1c6e8c34420b7630e3b06bc509d). The data will be made public upon publication of the manuscript.

## Code Availability

Relevant code repositories are referenced throughout the Methods sections.

## Supplementary Materials

Tables S1 to S11

## Supplemental figures

**Figure S1.**
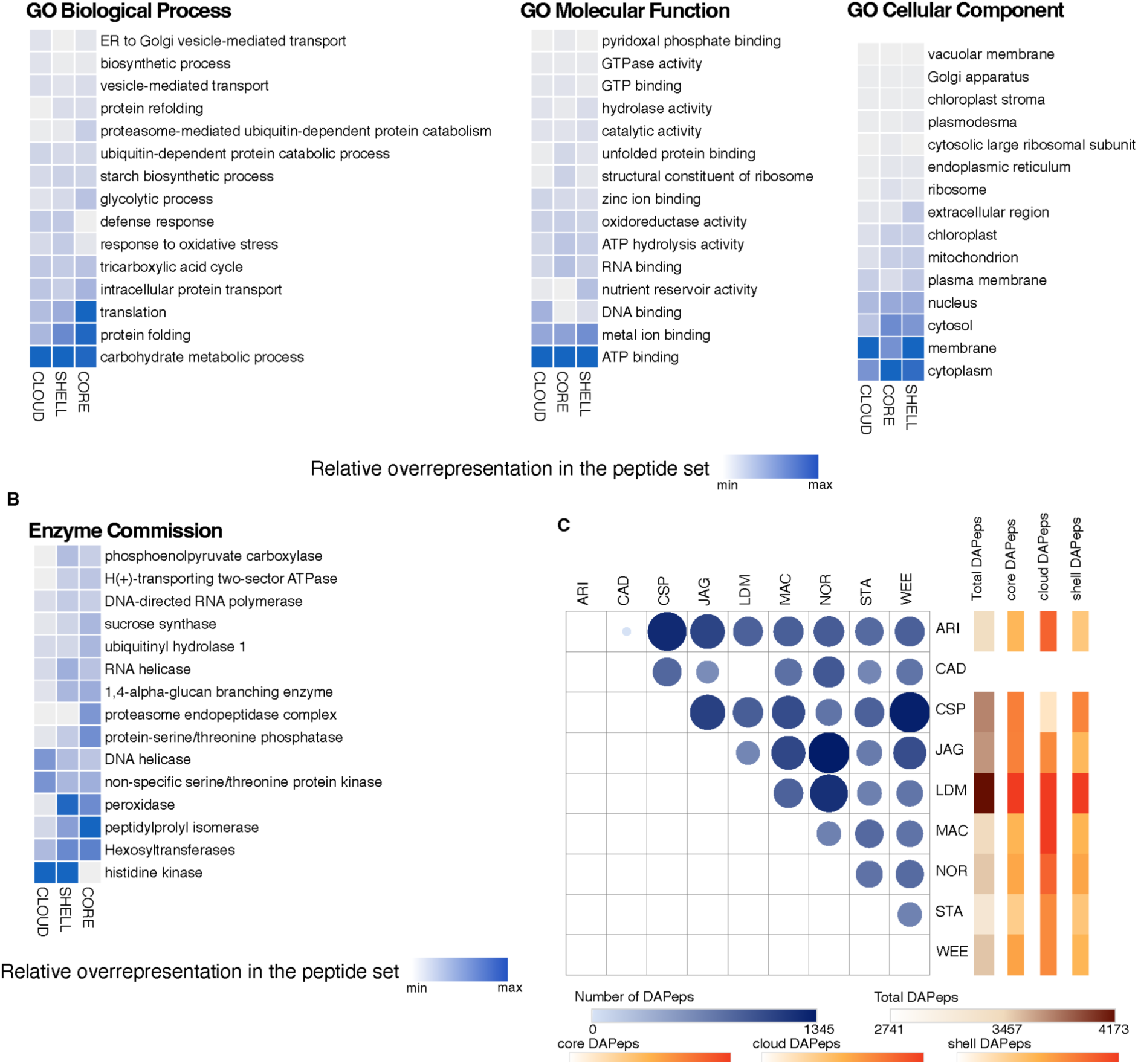
Comparison of the core, shell and cloud peptidomes. (**A**) Peptides detected in each cultivar (core), in at least two cultivars (shell) and characteristic of a single cultivar (cloud) were used for functional overrepresentation analysis in Uniept v. 2.0.2. The top 15 GO Biological Process, GO Molecular Function and GO Cellular Component. (**B**) Enzyme Commission classes are visualised on a light blue dark blue scale. C: Comparison of number of differentially abundant peptides (|log2FC|>1, adj. p-value <0.05. Abbreviation of the cultivars: ARI - Arina*LrFor*, CAD - Cadenza, CSP - Chinese Spring, JAG - Jagger, LDM - CDC Landmark, MAC – Mace, NOR – Norin 61, STA – CDC Stanley, WEE – Weebil.

**Figure S2.**
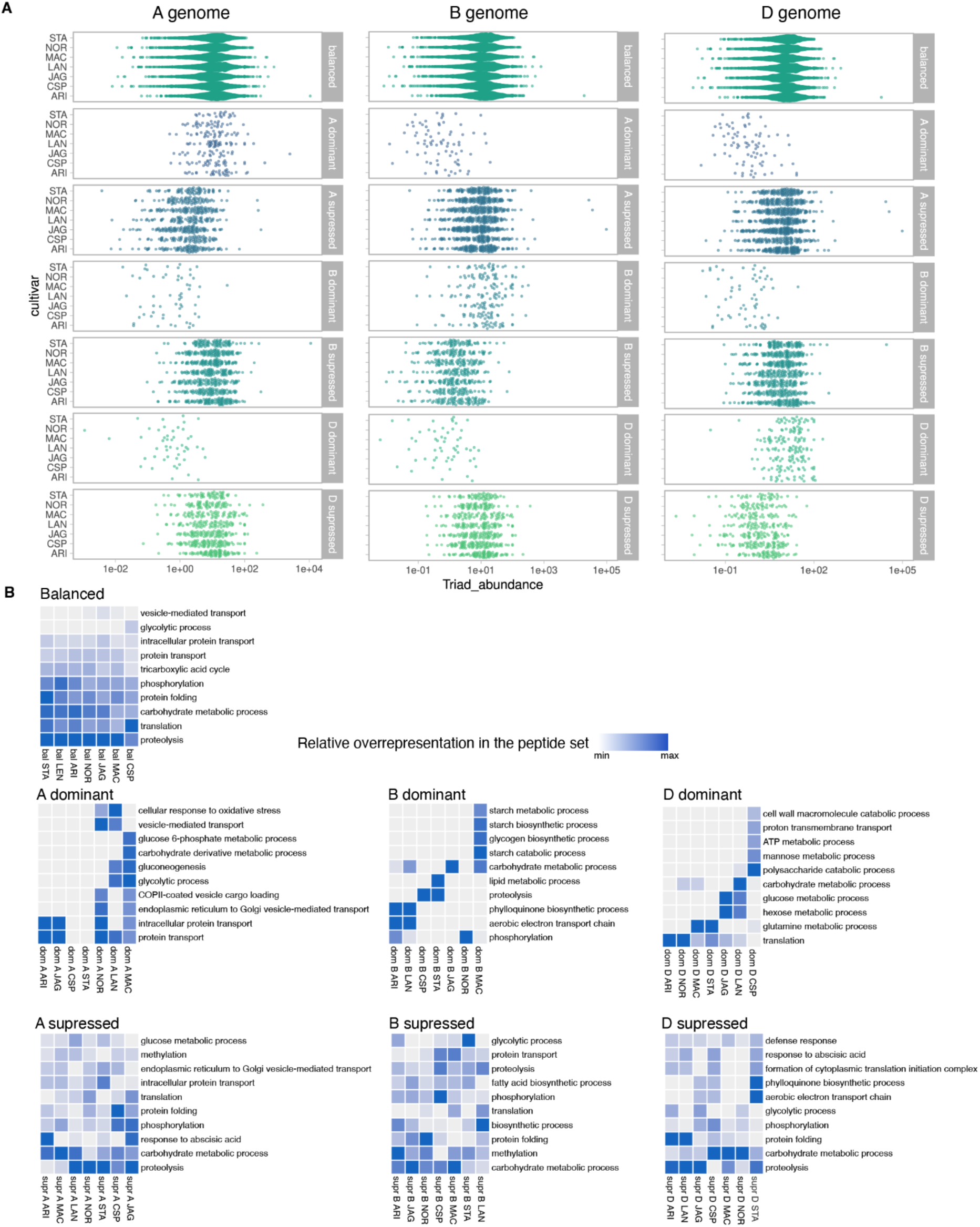
**Expression levels and functional characterisation of triad gene encoded proteoforms**. (**A**) Protein abundance of homeologous gene triad encoded proteoforms in the 7 chromosome assembled cultivars. (**B**) Overrepresented GO Biological process terms in the balanced, dominant and suppressed proteoforms.

**Figure S3.**
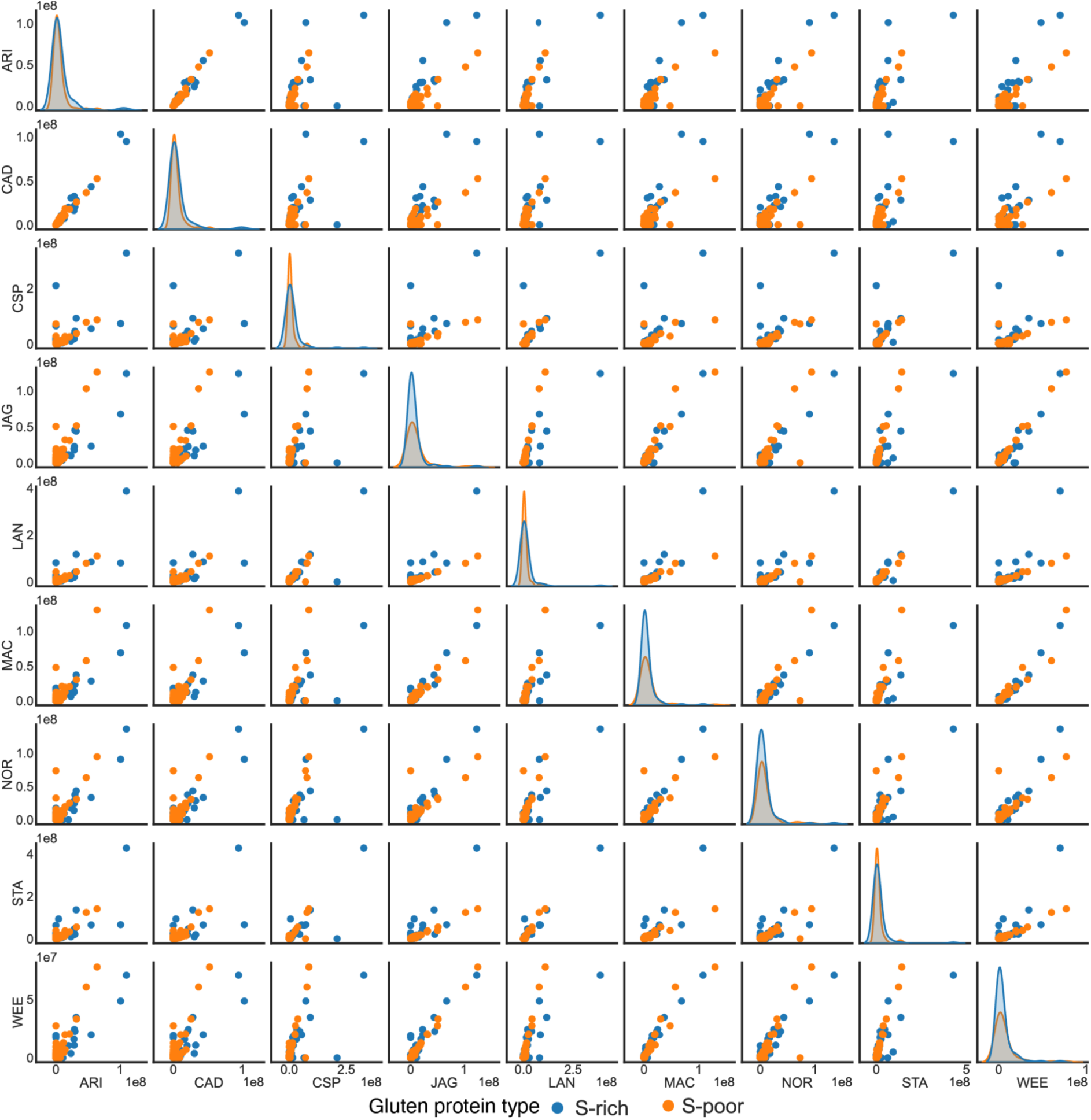
Comparison of S-rich and S-poor gluten protein profiles of the analysed cultivars. Targeted method was developed to assess differences in S-rich (alpha-, gamma-gliadin, LMW glutenin) and S-poor (omega-gliadin, HMW glutenin) gluten protein abundance values. Individual and total abundance values are visualised for each cultivar comparison.

**Figure S4.**
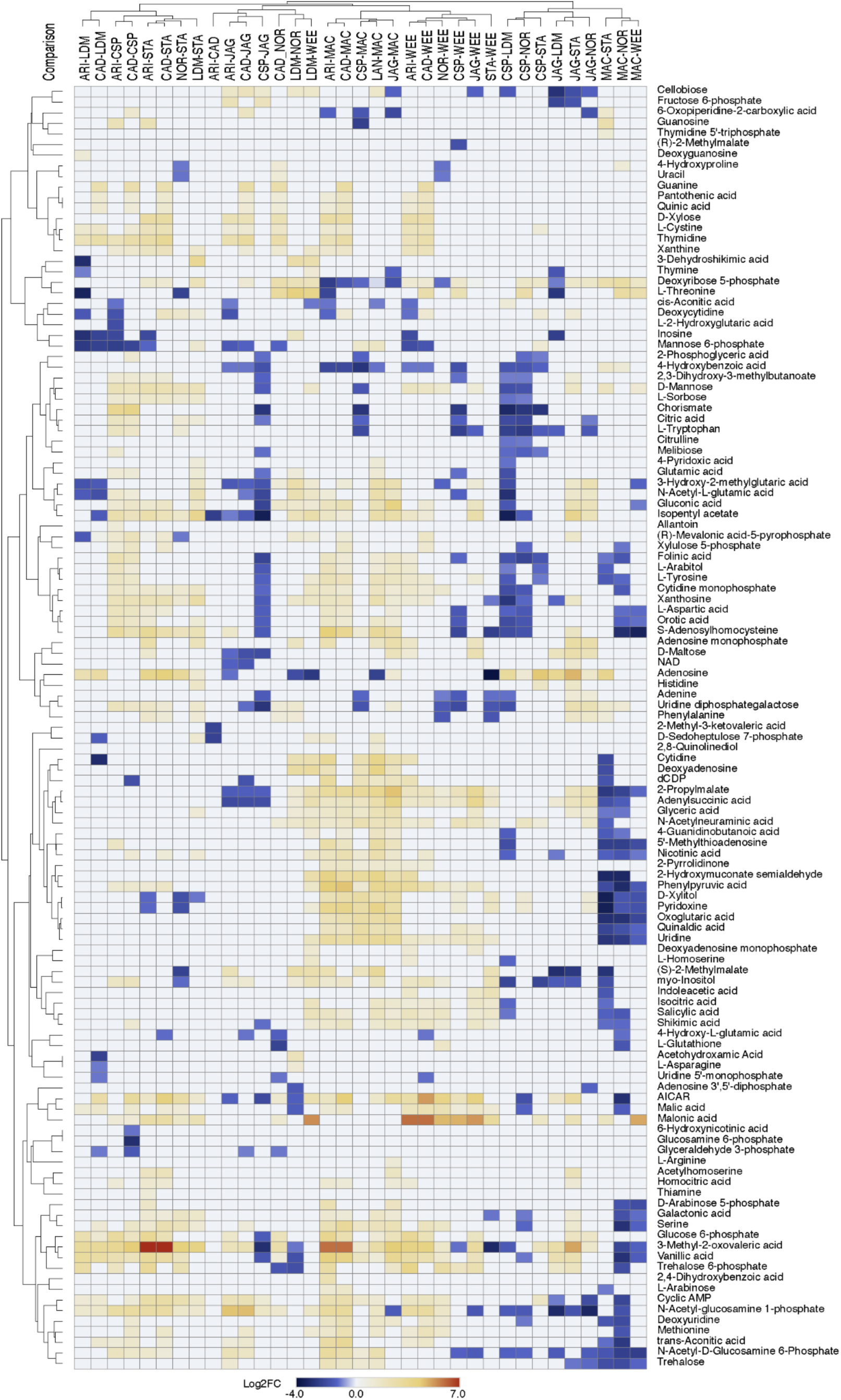
Comparison of differentially abundant metabolites across the cultivars. Significantly different metabolites with adj. p-value < 0.05 and |log2FC|>1 are visualized.

**Figure S5.**
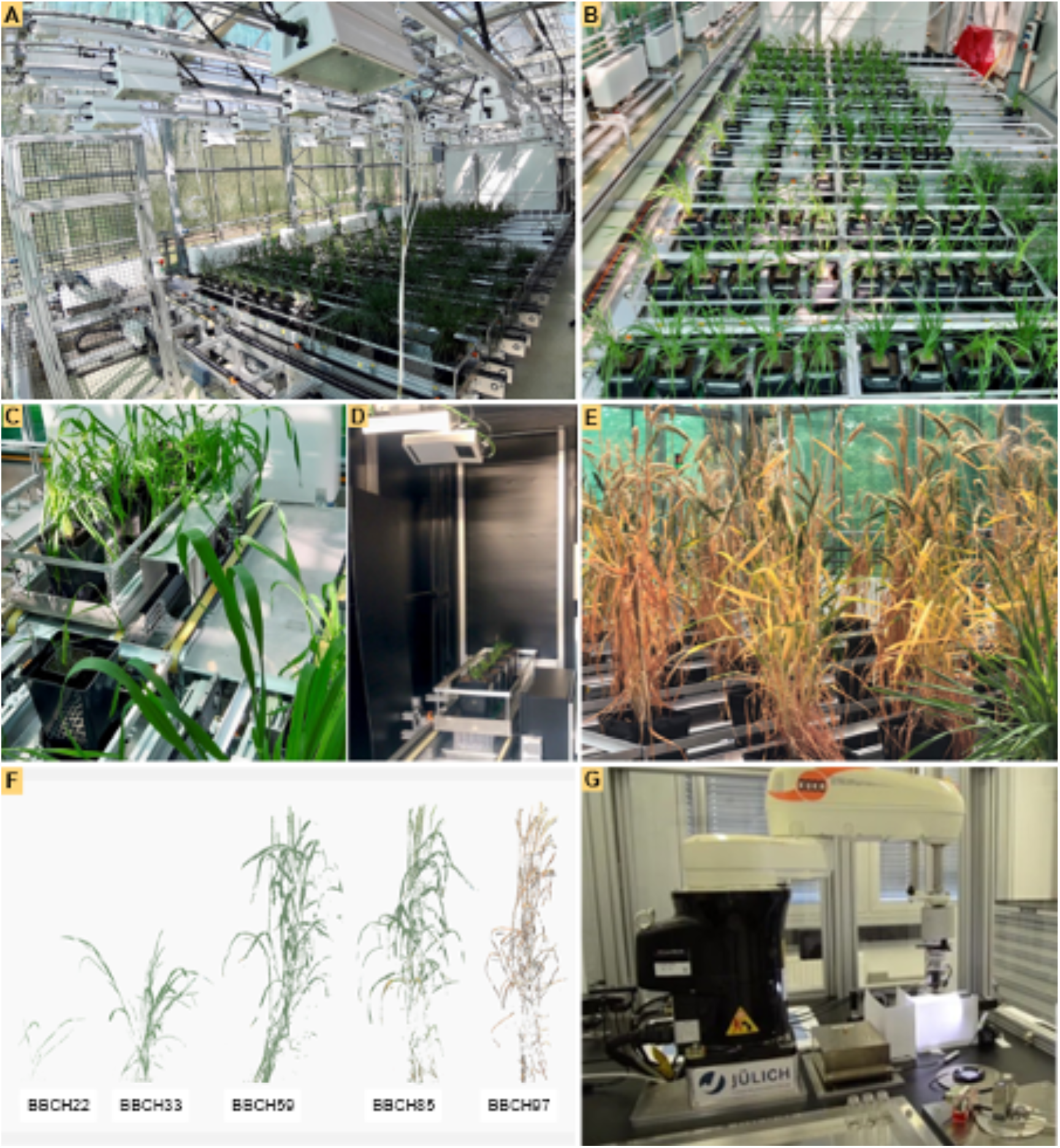
FitnessSCREEN phenotyping facility at Helmholtz Munich (HMGU) and seed phenotyping system at Forschungszentrum Jülich (FZJ). (**A**) conveyor belt transport system for transporting the pots to the imaging and watering stations. (**B**) pots in transport carriers, each carrier has a capacity of five pots. The number of pots was reduced to three plants per carrier when plants were overlapping in the scans. (**C**) carrier ready for transport to the imaging/scanning unit. (**D**) carrier in the imaging unit, the pot in front is positioned on the scales. (**E**) plants at BBCH stage 97. (**F**) true-to-life 3D representation of plants of the cultivar CDC Stanley at different developmental/BBCH stages. (**G**) “phenoSeeder” at FZJ.

**Figure S6.**
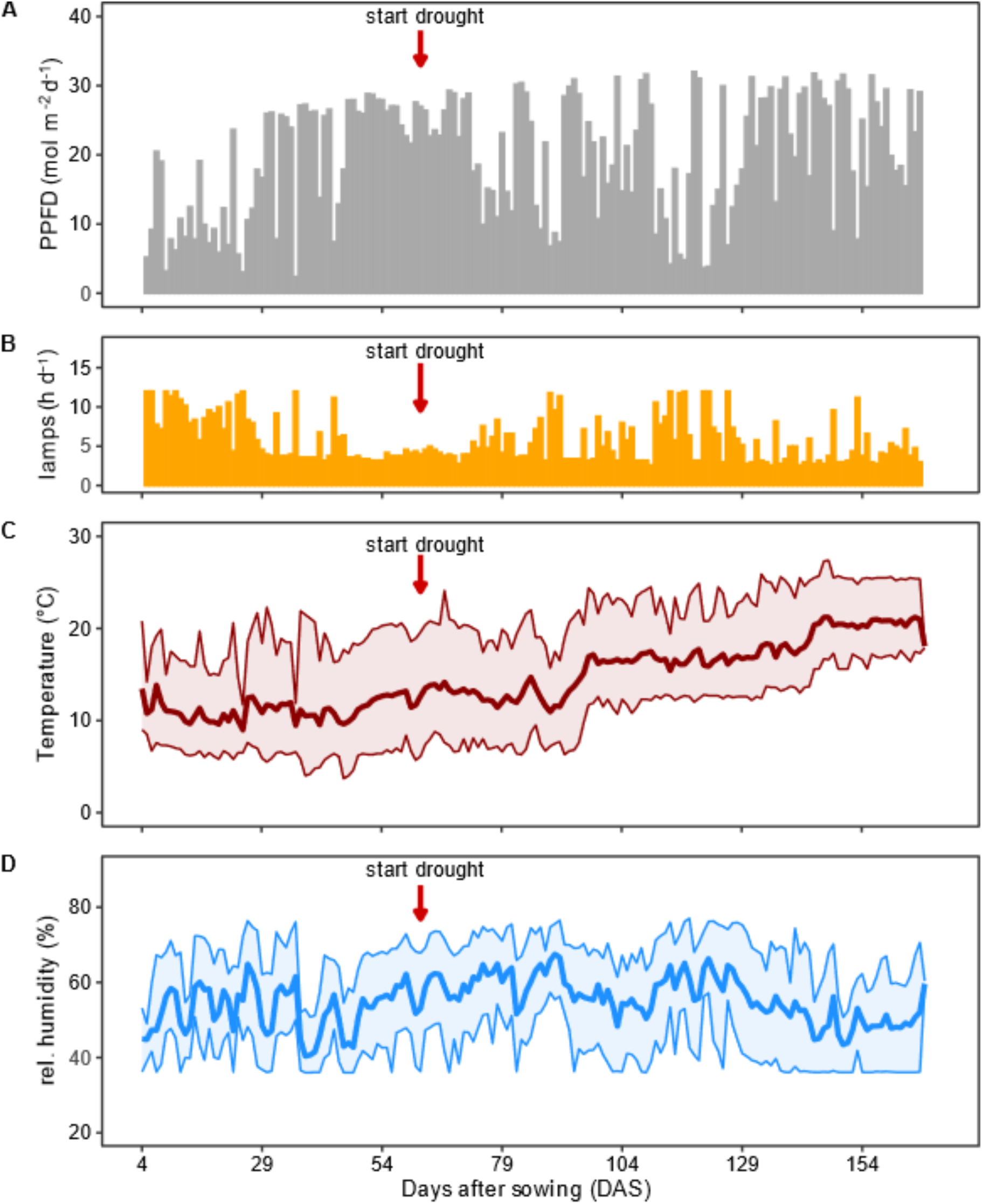
Ambient climatic conditions wheat cultivation on the FitnessSCREEN platform from day after sawing (DAS) 4 to 166. (**A**) Daily light integral (PPFD (mol m-2 d-1). Light was measured outside the greenhouse. (**B**) additional light by high pressure lamps, provided during daylight hours when outside light dropped to less than 370 µmol m-2 s-1 PPFD (20 klux). (**C**) Air temperature and (**D**) Relative air humidity, measured in the greenhouse compartment. Given are daily averages (bold lines), maximum and minimum values (line).

**Figure S7.**
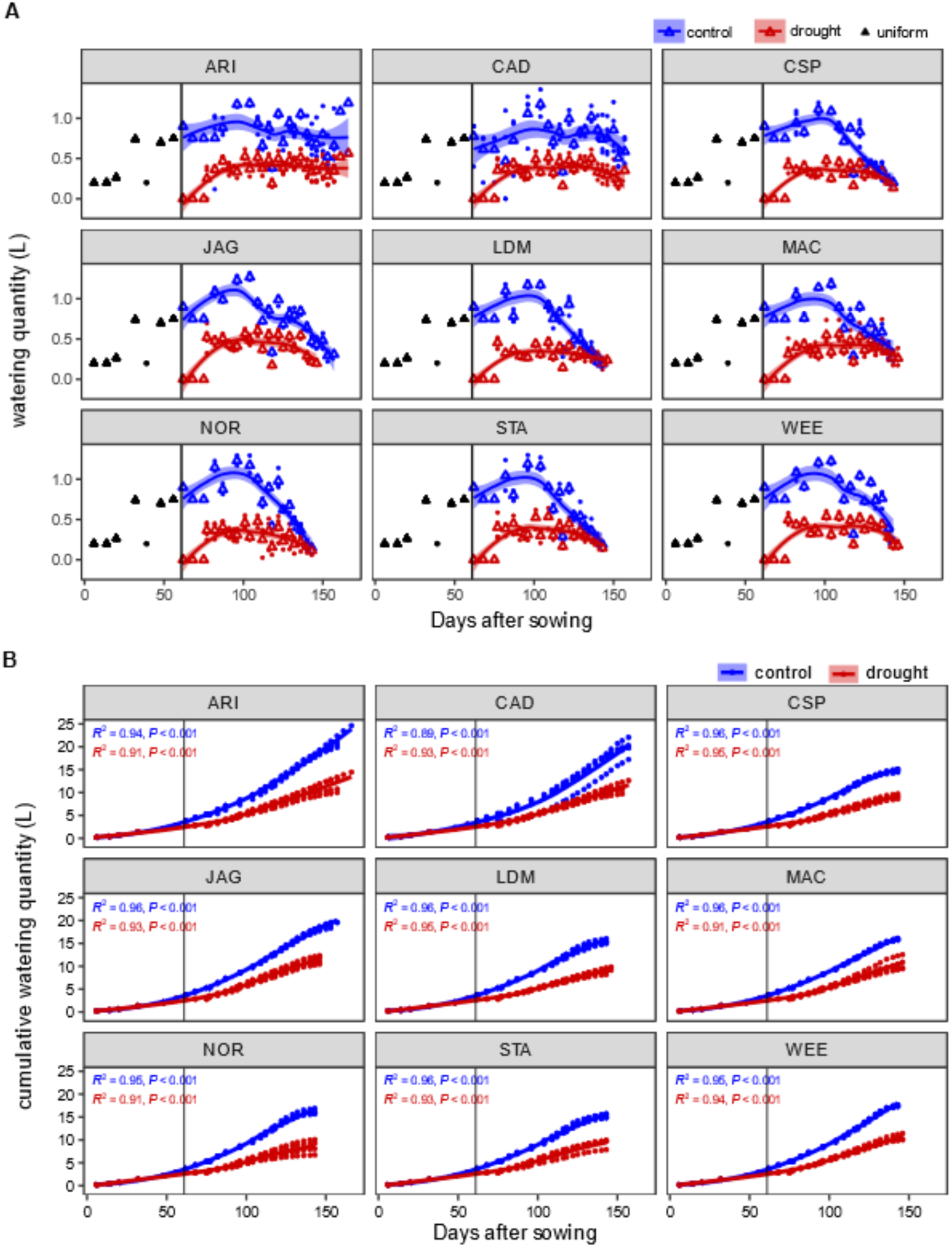
Irrigation regime during wheat cultivation on the FitnessSCREEN platform. (**A**) Watering quantity (L) per irrigation event. Until DAS 62, all plants received the same amount of water (black triangles) to keep 75% of “pot capacity” (water holding capacity of the pot). From DAS 62 onwards, plants received water according to pot weight (“on demand”), control plants (blue) were kept at 75% pot capacity, drought-treated plants (red) were watered up to 40% pot capacity (dots: values of individual plants, triangles: mean values per treatment). (**B**) Cumulative water supply for each plant (L) (blue: control irrigation, red: water limitation). Average per cultivar and irrigation level was fitted in **A** and **B** by a LOESS regression (line), quality of fitting is given by R^2^ (n=4, Arina*LrFor* n=3). The vertical line at DAS 62 marks the beginning of the drought stress treatment.

**Figure S8.**
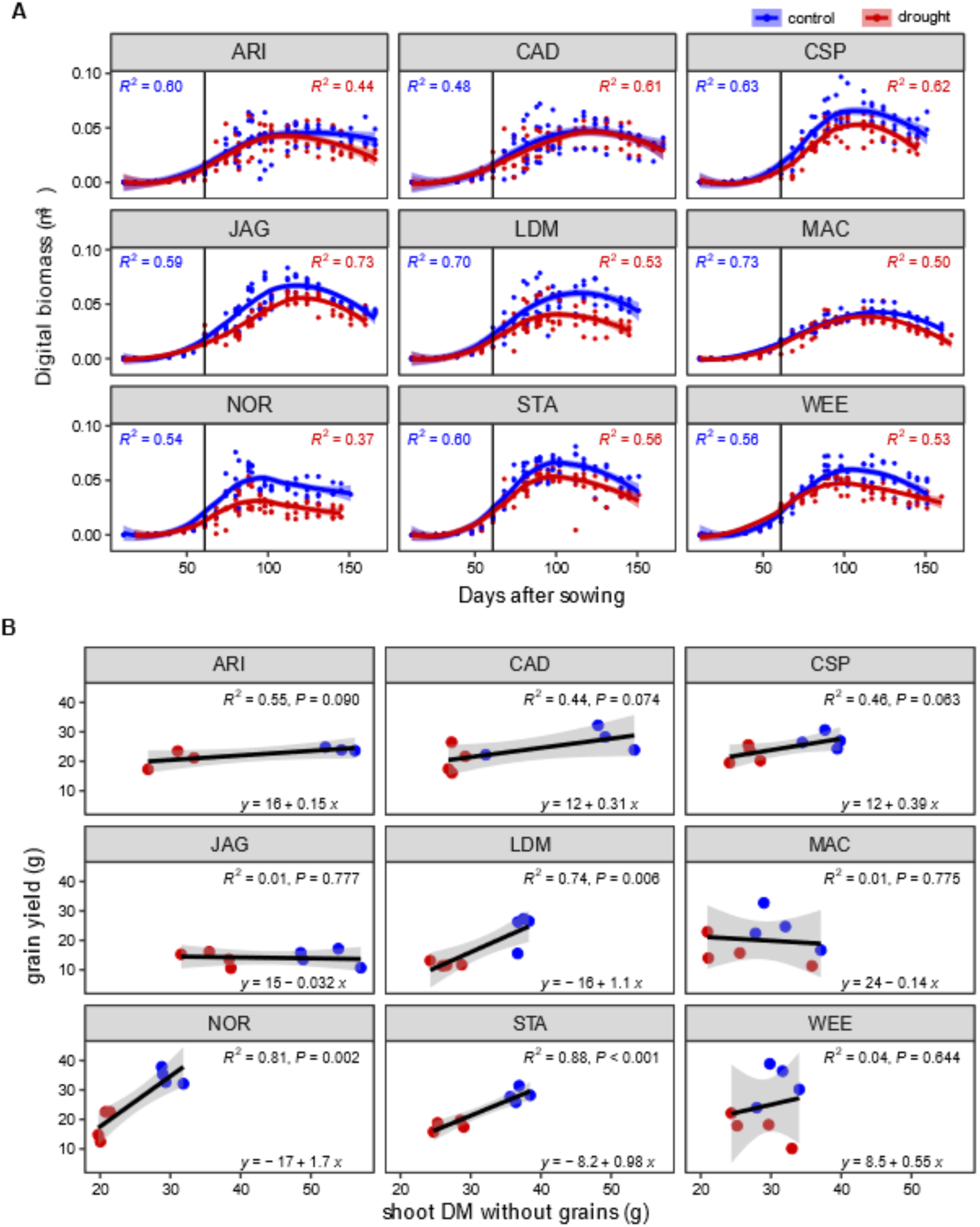
Plant morphological traits during cultivation on the FitnessSCREEN platform. (**A**) Digital biomass (m^3^) over the course of the experiment under control (blue) and water-limiting conditions (red). Data of individual plants (points, n=4, ARI n=3) and dynamics for each variety, fitted using LOESS regression (line), are depicted for the respective irrigation regime (the quality of the fit is indicated by R^2^). The vertical line at DAS 62 marks the beginning of the drought stress treatment. (**B**) linear model of grain yield and above-ground shoot dry mass (blue: control irrigation, red: water limitation, n=4, (Arina*LrFor* n=3)).

**Figure S9.**
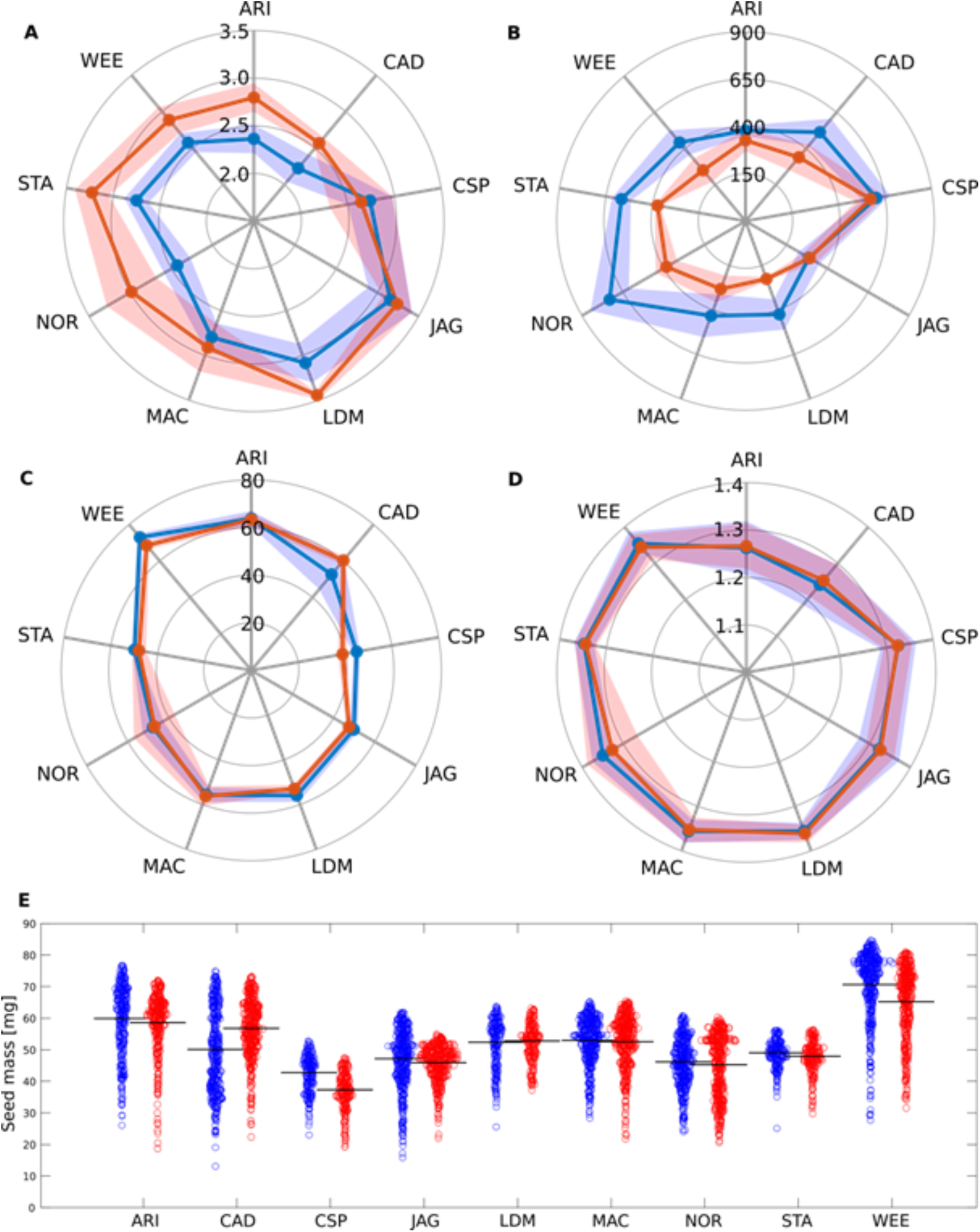
Phenotypic plasticity of wheat seeds developed under control and drought stress conditions. Selected traits of seeds from nine wheat cultivars grown under control (blue) and drought (red) conditions, respectively. (**A**) N content [%]. (**B**) Grain number per plant, calculated from total number of kernels and number of plants per cultivar. (**C**) 1000-grain weight [g]. (**D**) Mean seed density [mg/mm³]. (**E**) Individual seed mass. Shaded areas in (**A**-**D**) represent standard deviations.

**Figure S10.**
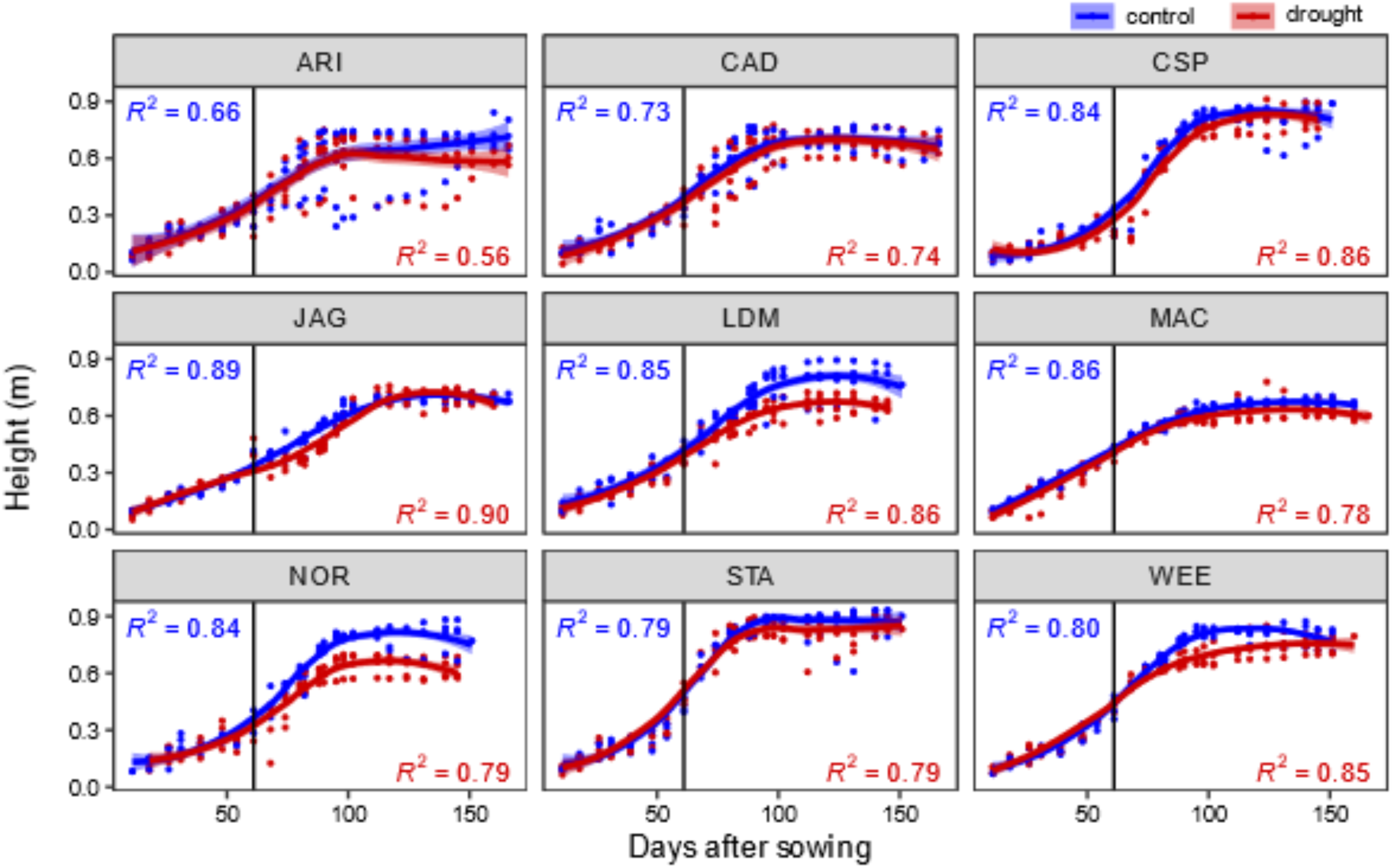
Development of plant heights during wheat cultivation on the FitnessSCREEN platform. Plant height (m) over the course of the experiment under control (blue) and water-limiting conditions (red). Data of individual plants (points, n=4) and dynamics for each variety, fitted using LOESS regression (line), are depicted for the respective irrigation regime (the quality of the fit is indicated by R^2^). The vertical line at DAS 62 marks the beginning of the drought stress treatment.

**Figure S11.**
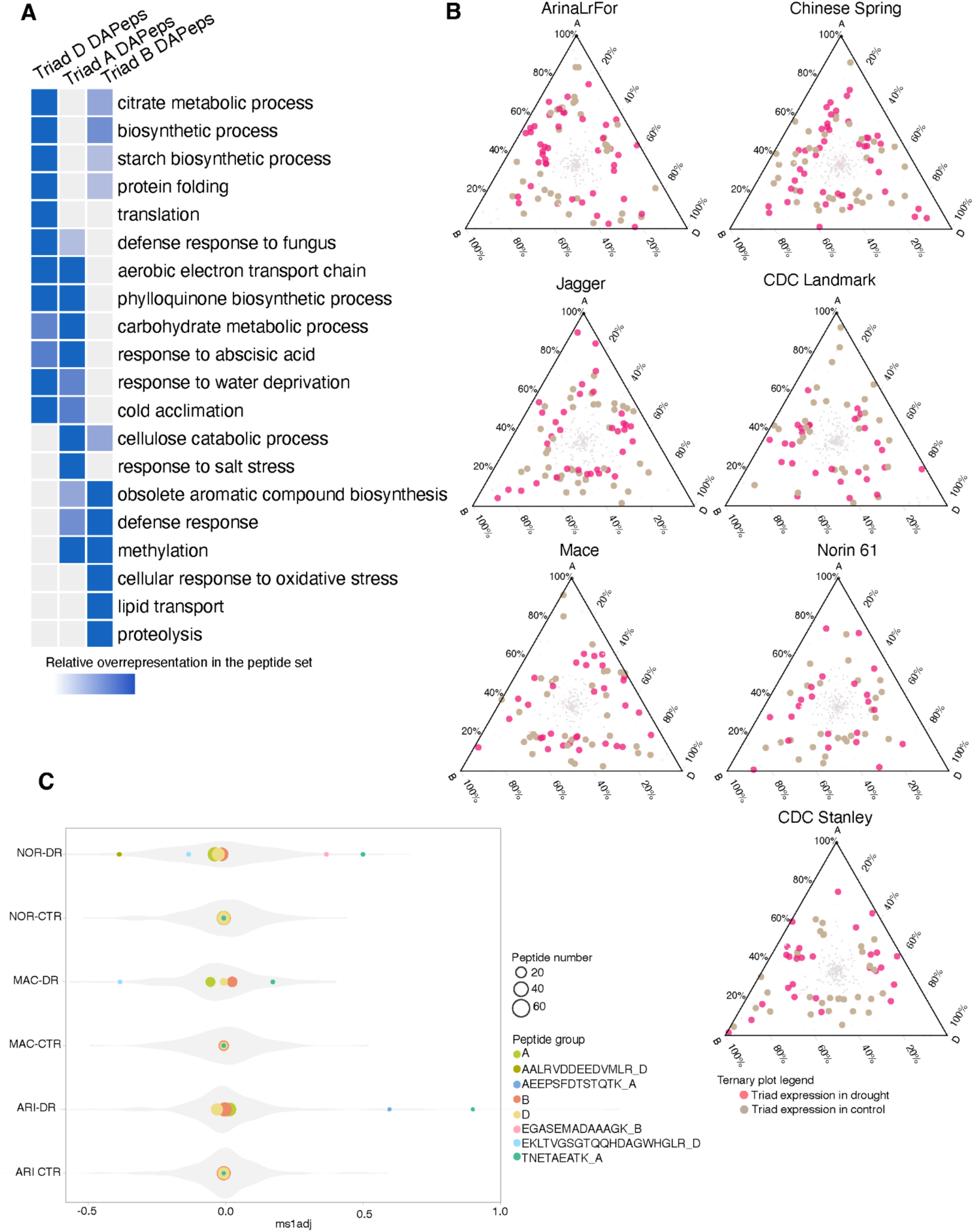
Impact of drought stress on triad proteoform levels. (**A**) Overrepresentation of GO Biological Process terms in the differentially abundant A, B and D subgenome triad proteins. (**B**) Ternary plots showing triad proteoforms with a minimum of 50% change in their abundance levels between control and drought condition. (**C**) Peptide abundance biases in chromosome 1 LEA proteoforms showing differentially regulated peptides. Peptides positioned at ms1adj value 0 represent balanced expression, peptides shifted towards the minimum and maximum value show biased levels.

**Figure S12.**
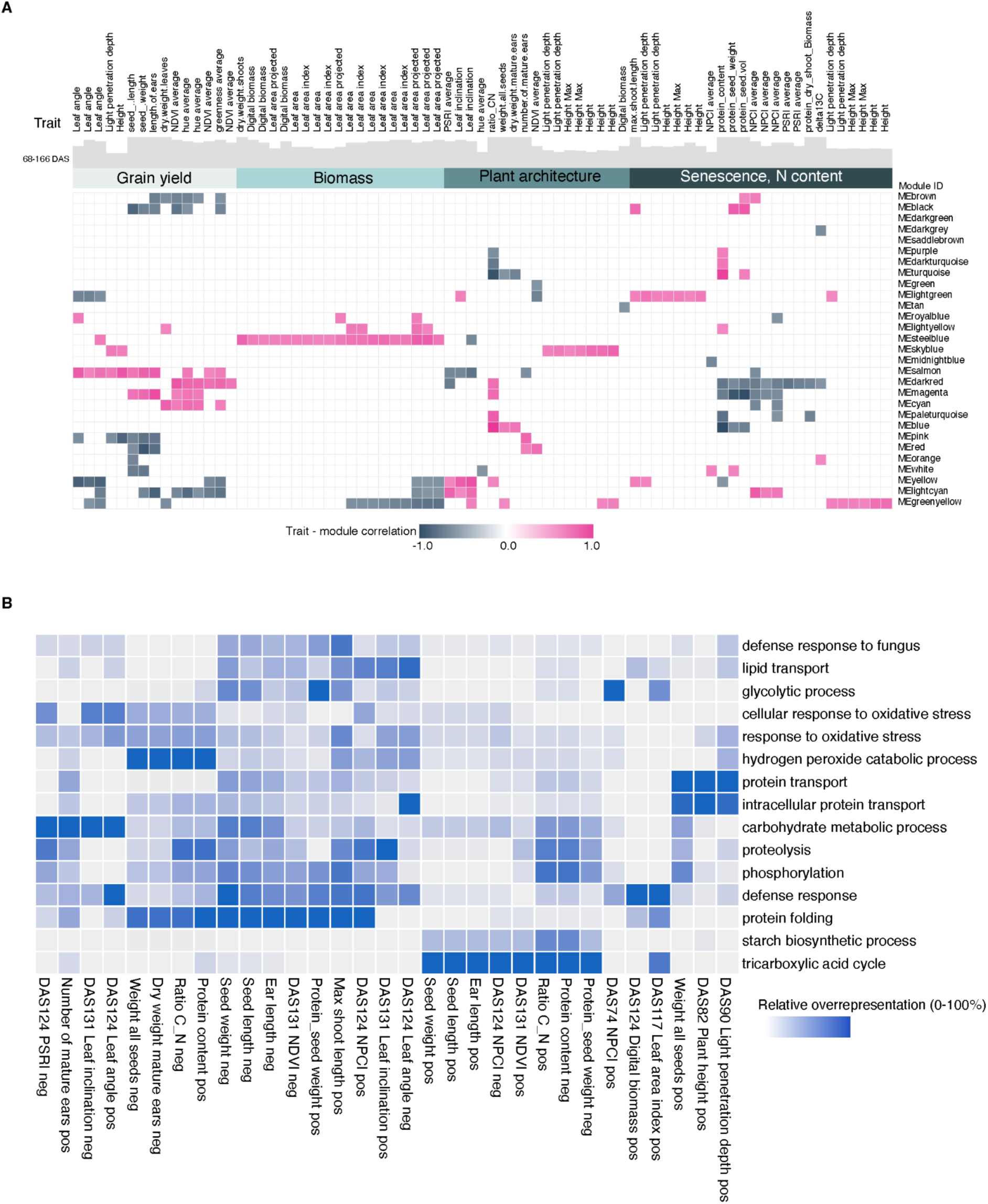
Trait - module relationship analysis of quantified proteins and measured plant phenotypes. (**A**) WGCNA analysis was used to define the protein modules and relate them to phenotype traits that show significant differences between control and drought samples during the lifecycle of the plants. Significant trait module relationships (adjusted p-value <0.05) are visualised. (**B**) GO Biological process overrepresentation analysis of protein modules with significant positive or negative relationship with the measured traits.

**Figure S13.**
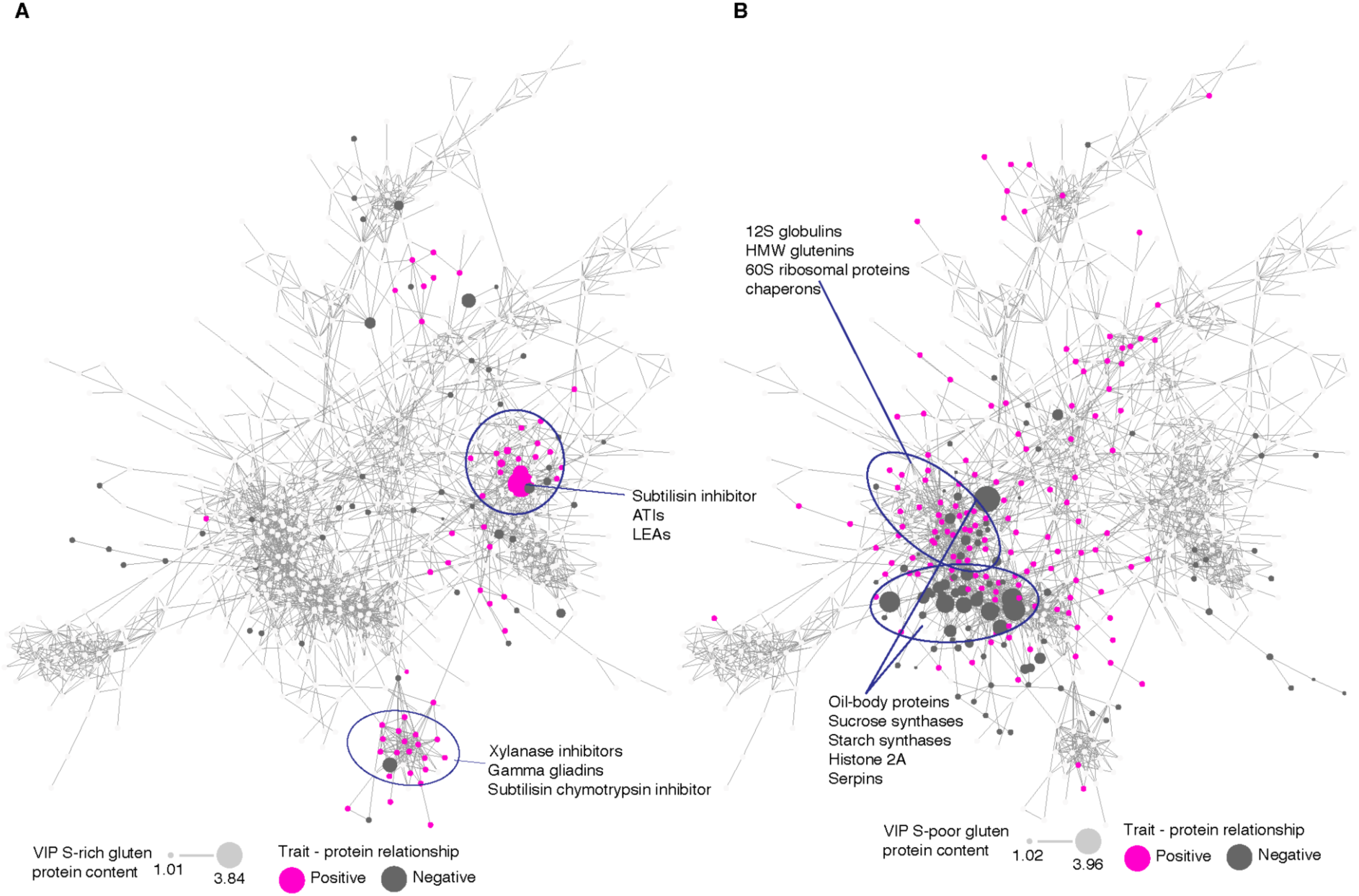
Protein co-abundance network analysis overlaid with key proteins associated with selected traits. Protein network was created using Pearson correlation cut-off value 0.8. Proteins positively or negatively associated with the selected traits are highlighted in magenta and dark grey. Dot size is proportional to the VIP value identified using PLS-DA VIP analysis are visualised (adjusted p-value<0.05, VIP >1). (**A**) S-rich gluten protein content related are highlighted. (**B**) S-poor gluten protein content related proteins.

**Figure S14.**
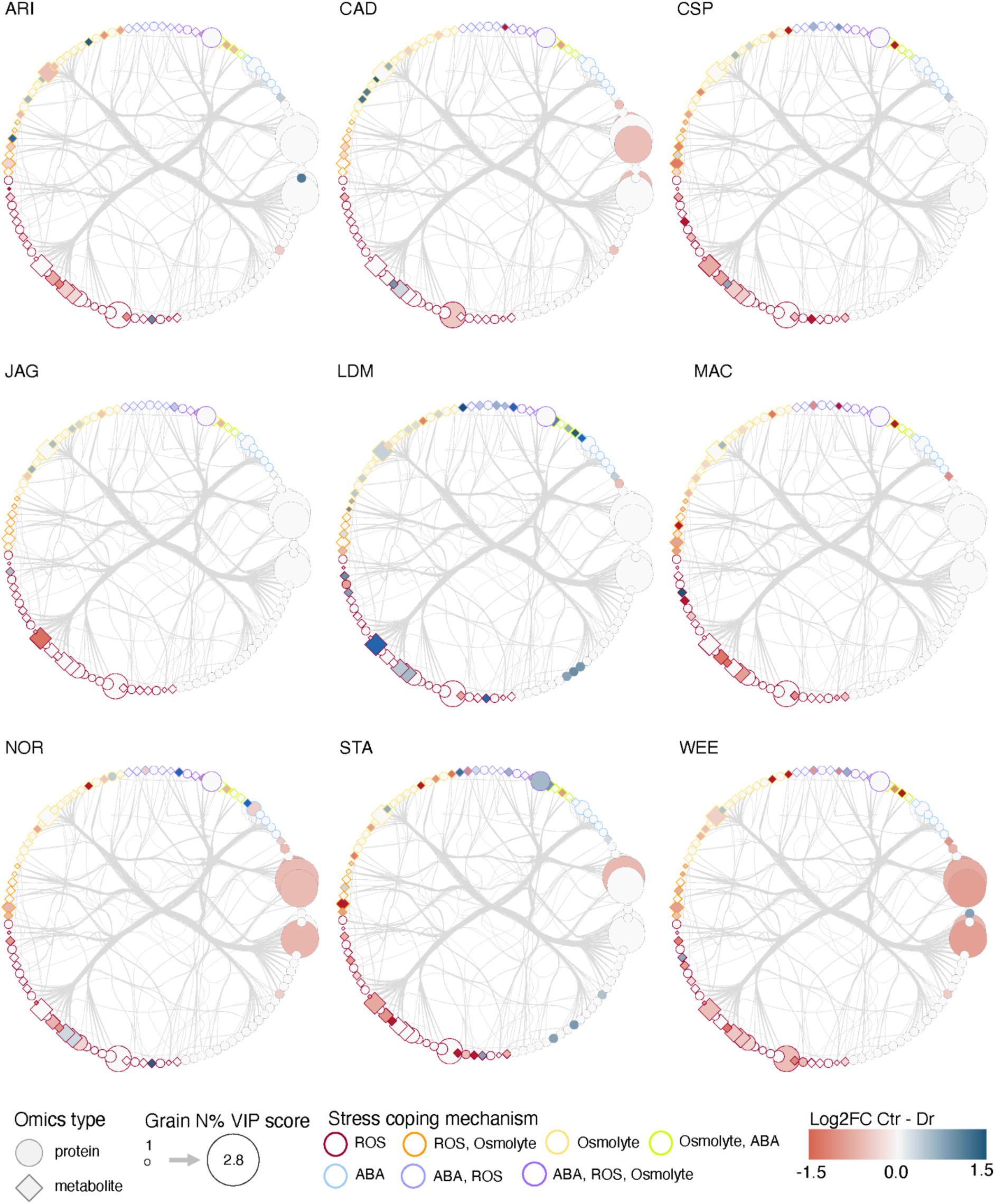
Control-drought fold change overlayed integrated protein and metabolite network. Proteins and metabolites filtered using DIABLO are visualised. Fold change between control and drought conditions are visualised on a red-blue scale with red colours representing increased abundance levels in drought condition. Feature sizes are proportional to the VIP

## Notes

### Competing Interest Statement

The authors have declared no competing interest.

